# Shieldin and CST co-orchestrate DNA polymerase-dependent tailed-end joining reactions independently of 53BP1-governed repair pathway choice

**DOI:** 10.1101/2023.12.20.572534

**Authors:** Ashleigh King, Pia Reichl, Jean S. Metson, Robert Parker, Daniella Munro, Catarina Oliveira, Jordan R. Becker, Daniel Biggs, Chris Preece, Benjamin Davies, J. Ross Chapman

**Affiliations:** Genome Integrity laboratory, Medical Research Council Molecular Haematology Unit, MRC Weatherall Institute of Molecular Medicine, Radcliffe Department of Medicine, The University of Oxford, Oxford, UK; Wellcome Centre for Human Genetics, University of Oxford, Oxford, UK; Centre for ImmunoOncology, Nuffield Department of Medicine, University of Oxford, Oxford, UK; Francis Crick Institute, 1 Midland Rd, London, UK

## Abstract

53BP1 regulates DNA end-joining in lymphocytes, diversifying immune antigen receptors. This involves nucleosome-bound 53BP1 at DNA double-stranded breaks (DSBs) recruiting RIF1 and shieldin, a poorly understood DNA-binding complex. The 53BP1-RIF1-shieldin axis is pathological in *BRCA1*-mutated cancers, blocking homologous recombination (HR) and driving illegitimate non-homologous end-joining (NHEJ). However, how this axis regulates DNA end-joining and HR suppression remains unresolved.

We investigated shieldin and its interplay with CST, a complex recently implicated in 53BP1-dependent activities. Immunophenotypically, mice lacking shieldin or CST are equivalent, with class-switch recombination co-reliant on both complexes. ATM-dependent DNA damage signalling underpins this cooperation, inducing physical interactions between these complexes that reveal shieldin as a DSB-responsive CST adaptor. Furthermore, DNA polymerase ζ functions downstream of shieldin, establishing DNA fill-in synthesis as the physiological function of shieldin-CST. Lastly, 53BP1 suppresses HR and promotes NHEJ in BRCA1-deficient mice and cells independently of shieldin. These findings showcase the resilience of the 53BP1 pathway, achieved through the collaboration of chromatin-bound 53BP1 complexes and DNA end-processing effector proteins.

## INTRODUCTION

DNA double-strand breaks (DSBs) are highly toxic and must be repaired accurately to counteract the threat of human disease and oncogenic mutations^1^. Paradoxically, mutagenic DSB repair by non-homologous end joining (NHEJ) is favoured in developing and antigen-stimulated lymphocytes, where it mediates deletional recombination events that diversify the antigenic specificity and function of B and T cell receptors^2^. To cope with this intrinsic discrepancy in desired DNA repair outcome between different cellular contexts, cells have evolved complex regulatory systems that maintain an appropriate equilibrium between competing DNA repair pathways, and that ensure DNA breaks are appropriately resolved^3^.

The generation of functional B and T cell receptors by the mammalian adaptive immune system relies on two programmed gene rearrangement mechanisms: V(D)J recombination and Class-Switch Recombination (CSR)^2^. Both mechanisms rely on intragenic deletional recombination events, which are mediated through the ligation of interspaced DNA ends using non-homologous end-joining (NHEJ) pathways. Due to the distinct biochemical mechanisms involved in generating DNA double-strand breaks (DSBs) during V(D)J recombination and CSR, the resulting DNA ends have different structural properties, necessitating distinct DNA-end processing and ligation mechanisms^4^.

DSB induction during V(D)J recombination occurs during the development of B and T cells in the bone marrow and thymus, respectively. The process is orchestrated by Recombination Activating Gene enzymes 1 and 2 (RAG1/RAG2), which cleave V, D and J segment-flanking recombination signal sequences (RSSs) resulting in pairs of blunt signal and covalently sealed hair-pinned coding ends. Hairpin opening is essential before coding-joins can occur; a step facilitated by the Artemis endonuclease upon its activation by the DNA-dependent Protein Kinase (DNA-PK) complex. This generates DNA ends that are strictly joined by classical NHEJ (c-NHEJ) proteins including Ku (Ku70/Ku80), XRCC4 and DNA Ligase 4 (LIG4), with a more modest reliance on XLF^2^.

In contrast, class-switch recombination in mature peripheral lymphocytes involves DSB generation between antibody isotype-encoding constant (C) gene segments of the immunoglobulin heavy chain locus (*igh).* This process is initiated by Activation-Induced Cytidine Deaminase (AID), which introduces clusters of U:G mismatches in actively transcribed switch *(S)* introns. These mismatches are then processed by base excision repair (BER) and mismatch repair (MMR) machineries, ultimately leading to the conversion of single-stranded DNA (ssDNA) gaps into DSBs with single-stranded tailed termini. Unlike RAG1/2-induced DSBs, AID-dependent DSBs during CSR can be repaired through both c-NHEJ and alternative end-joining (a-EJ) pathways, with a critical role also played by the 53BP1 pathway proteins^4^.

The 53BP1 pathway is composed of nucleosomal and DNA-binding protein components. Most upstream is 53BP1, a chromatin reader protein scaffold that interacts with the DSB-responsive ubiquitinated Lysine 15 on nucleosomal H2A type histones (H2AK15ub) and ubiquitous histone H4 methylation at Lysine 20 (H4K20me1/2)^5,6^. This interaction leads to the formation of extensive 53BP1 chromatin-bound domains at DSB sites. Downstream of 53BP1, the 53BP1-binding protein RIF1 bridges interactions between ATM-phosphorylated motifs in 53BP1 and the SHLD3 subunit of shieldin^7,8^, a complex bound to single-stranded DNA (ssDNA) that is thought to protect DNA ends^9^. Studies using genetically engineered mouse models (GEMMs) deleted of *53bp1*, or shieldin proteins *Rev7* or *Shld1/2* present with near-complete deficiencies in CSR. By contrast, only *53bp1^-/-^* mice exhibit V(D)J recombination defects, which lead to mild B and T cell lymphopenia^10–13^.

The significance of the 53BP1 pathway extends to *BRCA1*-mutated cancers. In normal cells, the BRCA1-BARD1 heterodimer binds to a related combinatorial chromatin signature comprising H2AK15ub and unmethylated Lysine 20 on histone H4 (a post-replicative modification state), excluding 53BP1 from DNA damage sites and directing HR^14,15^. However, in *BRCA1-*deficient cells, uninhibited 53BP1-chromatin complexes prevent HR and promote NHEJ-mediated genome rearrangements, leading to genomic instability^16,17^. Consequently, *BRCA1*-deficient cells and tumours are hypersensitive to poly(ADP-ribose) polymerase inhibitors (PARPi), which exploit the HR defect caused by 53BP1^16^. Conversely, 53BP1-deletion is able to restore PARPi resistance to *BRCA1-*mutant cells^18^, and can even rescue viability in *Brca1-*null mice^19,20^.

Genetic screens aimed at elucidating PARPi resistance mechanisms in *BRCA1*-deficient cells were instrumental in revealing 53BP1 pathway biology: shieldin genes were discovered through screens investigating PARPi-induced toxicity in BRCA1-deficient cells^21–23^; the Polɑ-Primase accessory complex proteins Ctc1-Stn1-Ten1 (CST) similarly emerged as putative 53BP1 pathway effectors^24^. Concurrent hypothesis-guided studies also linked the activities of shieldin and CST to pathological 53BP1 pathway function in *BRCA1-*mutant cells^11,25–28^, and highlighted CST-directed Polɑ-Primase-dependent DNA fill-in synthesis as key intermediary step^27,29,30^.

This research has ignited debates regarding the precise role of shieldin, particularly whether its primary function involves blocking nucleolytic resection or facilitating fill-in synthesis at DNA ends^4,31^. The importance of effector interactions in the 53BP1 pathway during physiological antigen receptor recombination mechanisms, as well as their impact on HR defects and DNA joining events in BRCA1-deficient cells, remains to be fully understood. To address these questions, we conducted a comprehensive genetic analysis using GEMMs. Our findings demonstrate that the 53BP1 pathway contributes substantially to DNA end-joining during antigen receptor recombination and in cancer cells in the absence of shieldin/CST. We also demonstrate a strict interdependence between shieldin and CST during CSR, and an epistatic relationship with DNA polymerase ζ, solidifying fill-in synthesis as a primary shieldin-dependent DNA repair function. Importantly, using *BRCA1-*deficient mice and cellular models we establish that HR suppression relies on 53BP1 but not shieldin. Altogether, our results showcase remarkable resilience in the 53BP1 pathway, achieved through a division of labour between chromatin-bound 53BP1 complexes and its downstream DNA-bound effectors.

## RESULTS

### Shieldin and CST equivalency during lymphocyte development

To understand the extent to which shieldin-CST cooperation supports physiological 53BP1 pathway function, we generated new mouse strains deleted for genes encoding core proteins in each complex in an inbred C57BL/6 background. Knockout mice deleted for critical exons 4-5 in *Shld2,* or exon 2 in *Shld3* which comprises its entire coding sequences (hereafter referred to as *Shld2*^-/-^ and *Shld3*^-/-^ mice; (Fig. 1A; Extended Data Fig. 1A and 1B) were viable, healthy, fertile and born at expected mendelian frequencies. Moreover, genomic instability in 8–12-week-old *Shld2*^-/-^ and *Shld3*^-/-^ mice did not exceed that seen in aged-matched cohorts of wild-type or *53bp1^-/-^* mice, as defined by levels of micronuclei in erythrocytes (Extended Data Fig. 1D and 1E), a proxy for systemic genomic instability^32^.

**Fig. 1:**
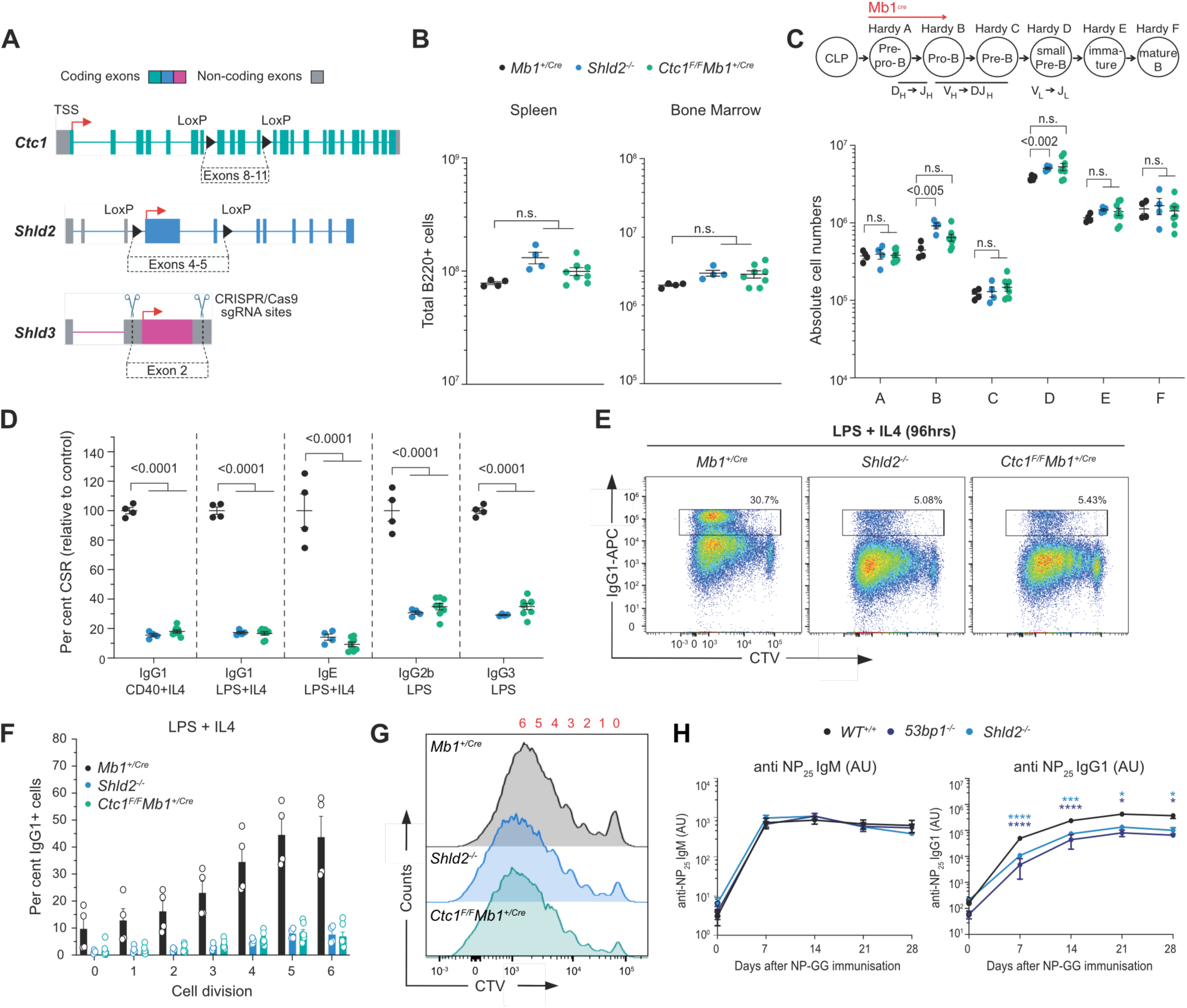
Equivalent immunodeficiencies in mice lacking shieldin and CST. **A)** Schematic representation of the *Shld2* and *Shld3* (knockout), and *Ctc1* (conditional) alleles generated in this study. **B)** Absolute numbers of B220+ B cells in the bone marrow (one femur and one tibia) and spleen. n = 4-8 mice per genotype. Significance was determined by unpaired two tailed *t*-test, Mean ± SEM. **C)** Absolute numbers of B cell precursors (Hardy fraction A, B220^+^CD43^+^BP-1^−^CD24^−^; Hardy fraction B, B220^+^CD43^+^BP-1^−^CD24^+^; Hardy fraction C, B220^+^CD43^+^BP-1^+^CD24^+^; Hardy fraction D, B220^+^CD43^−^IgM^−^IgD^−^; Hardy fraction E, B220^+^CD43^−^IgM^+^IgD^−^; and Hardy fraction F, B220^+^CD43^−^IgM^+^IgD^+^) in the bone marrow (one femur and one tibia). n = 4-8 mice per genotype. Significance was determined by unpaired two tailed *t*-test, Mean±SEM. **D)** Splenic B cells cultured with the indicated stimuli (96 h) and stained for surface IgG1, IgE, IgG2b or IgG3. *n* = 4-7 mice per genotype. CSR 100%, mean immunoglobulin isotype switch frequency of 2 control animals in each experiment. Significance was determined by two-way ANOVA with Tukey’s correction. Mean ± SEM. **E)** Cell trace violet (CTV)-labelled splenic B cells were stimulated as indicated and stained for surface IgG1 after 96 h. Representative data, *n* > 6 mice. **F)** IgG1-positive (IgG1^+^) B cells as a proportion of total B cells (%) for each cell generation as determined by CTV staining and proliferation-associated dye dilution. n = 4-6 mice per genotype. Mean ± SEM. **G)** CTV dilution in purified B cells cultured in the presence of LPS and IL-4 for 96 h. Representative data, *n* > 6 mice. **H)** NP-specific serum IgM (left) and IgG1 (right) at indicated times after NP-CGG immunization. AU, arbitrary units. Representative data, n = 2 independent experiments, each with 4 mice. Mean ± SEM.

Mutations affecting the CST gene *CTC1* in humans cause Coats Plus, an autosomal recessive disorder characterized by retinal telangiectasia, intracranial calcifications, osteopenia, gastrointestinal bleeding, and in severe cases, normocytic anaemia reflecting a degree of BM failure^33^. Similarly, haematopoietic stem cell (HSC) exhaustion characterises mice homozygous for *Ctc1* gene deletions, leading to complete bone marrow failure and perinatal lethality^34^. International mouse phenotyping consortium (IMPC)^35^ phenotyping of mice homozygous for a knockout-first conditional (ready) allele in *Ctc1 (Ctc1^tm1a(KOMP)Wtsi^*) also discovered perinatal lethality in *Ctc1-*deleted mice^36^. *Ctc1* essentiality in murine haematopoiesis thus prompted us to investigate the effect of *Ctc1-*deletion in the B lymphocyte lineage, where the 53BP1 pathway supports normal development and programmed genome diversification. To this end, we generated conditional *Ctc1^tm1c/tm1c^ Mb1*^+/Cre^ mice (hereafter referred to as *Ctc1^F/F^ Mb1*^+/Cre^ mice), in which *Cre* expression from the *Mb1-Cre* transgene mediated complete deletion of *Ctc1* critical exons 8-11 in early B cell progenitors (Fig. 1A; Extended Data Fig. 1C). To our surprise, CST-deficiency was well-tolerated in developing and differentiating B lymphocytes: *Ctc1^F/F^ Mb1*^+/Cre^ mice showed normal cell frequencies across all stages of lineage development in the bone marrow, and B cell maturation in the spleen (Fig. 1B, 1C; Extended Data Fig. 1F, 1L and 1M). The normal development of *Ctc1^Δ/Δ^* B cells aligned with the phenotypes of *Shld2*^-/-^ and *Shld3*^-/-^ mice, which similarly fostered normal B and T cell lineage development (Extended Data Fig. 1G-K and 1O-P). Normal lymphocyte development in *Shieldin-*knockout mice owes to a proficiency in V(D)J recombination, as demonstrated in our previous characterisation of mice deleted of the shieldin gene *Rev7*^11^, and more recent concordant results in *Shld1* and *Shld2* mutant mice^12,13^. Thus, neither CST, nor shieldin contribute to lymphocyte development, where 53BP1 supports efficient DNA end-joining during V(D)J recombination^10,11^.

### Shieldin-CST cooperation underpins productive CSR

We next sought to define the importance of shieldin-CST interplay for the joining of AID-dependent DSBs during class-switch recombination (CSR), a process entirely dependent on 53BP1 that also requires the *Rev7* and shieldin proteins^11–13^. To this end, we stimulated cultures of mature B splenocytes from *Ctc1^F/F^ Mb1*^+/Cre^, *Shld2*^-/-^ and *Shld3*^-/-^ mice and analysed their ability to support antibody isotype switching *in vitro.* Strikingly, *ex-vivo* stimulation of mature splenic B cells from *Ctc1^F/F^ Mb1*^+/Cre^ mice revealed severe, >5-fold, defects in CSR across all analysed IG isotypes, fully recapitulating the magnitude of defect presented in *Shld2-*deficient B cells analysed in parallel (Fig. 1D and 1E). Defective class-switching in *Shld2* and *Ctc1*-deficient B cells did in no case correlate to defects or differences in cell proliferation or survival (Fig. 1F-G). Despite this, B cells from *Ctc1^F/F^ Mb1*^+/Cre^ mice and both *shieldin* knockout mouse strains supported higher class-switching frequencies than those from *53bp1^-/-^* mice, where CSR was reduced >10-fold relative to wild-type (Fig. 1D, 1E; Extended Data Fig. 1Q, and 1R). These differences also held true *in vivo,* in mice immunised with the model antigen NP-CGG (3-hydroxy-nitrophenyl acetyl coupled to chicken γ-globulin): following immunisation, serum titres of NP-specific IgG1 were strongly attenuated in both *Shld2^-/-^* and *53bp1^-/-^* mice relative to wild type controls, despite exhibiting wild type IgM-mediated antigen responses as expected (Fig. 1H); however, antigen-specific IgG1 consistently accumulated to higher levels in *Shld2^-/-^* mice than in *53bp1^-/-^* mice at all time-points following immunisation (Fig. 1H). Considered together our results show the 53BP1 pathway retains some capacity to support CSR in the absence of shieldin and CST. The fact that B lymphocyte development and differentiation capacities were found to be identical in *Ctc1^F/F^ Mb1*^+/Cre^ and *Shld2/3*-deficient mice, prompted us to next investigate the mechanistic basis of shieldin’s coordination with CST and downstream factors in CSR.

### DNA damage signalling supports shieldin-CST interplay during DSB repair

To investigate shieldin adaptor functions in DSB repair, we used proteomics to define the shieldin interactome. For this, we introduced TwinStrep-tagged *Shld1* and *Shld3* transgenes into *Shld1*^-/-^ and *Shld3*^-/-^ CH12-F3 mouse B cell lymphoma cell lines, respectively, and validated functional complementation at the level of rescued IgM to IgA class-switching in stimulated cell cultures (Extended Data Fig. 2A and 2B). Due to shieldin’s extreme low abundance in mammalian cells^26^, its successful purification required lysates prepared from cultures of > 6 x10^9^ cells. Shieldin complexes isolated via Streptactin affinity purification, were then subjected to analysis by liquid chromatography-tandem-mass-spectrometry (LC-MS/MS) over two independent experiments (Fig. 2A).

**Fig. 2:**
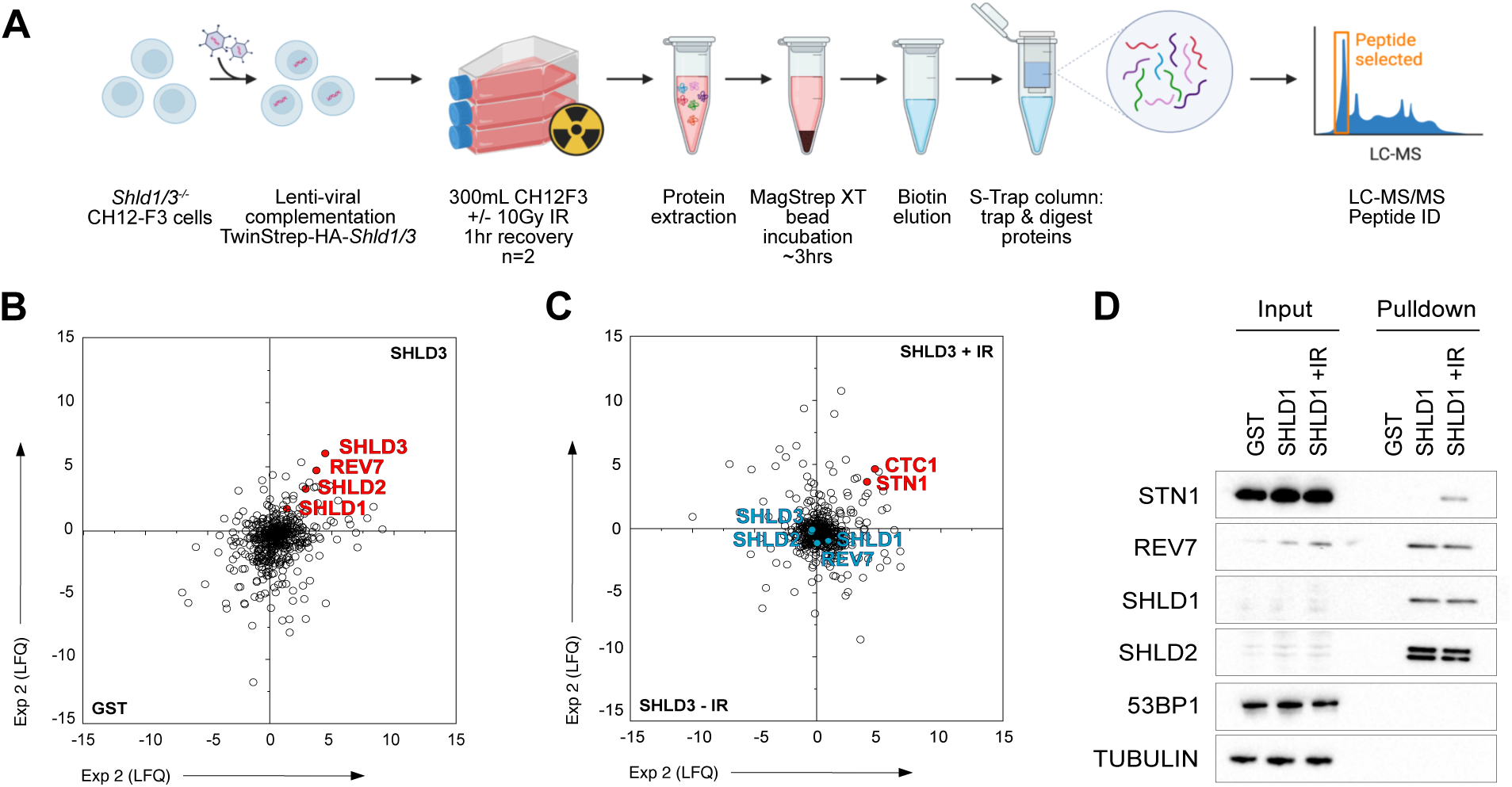
DNA damage induces interactions between shieldin and CST. **A)** Schematic depicting shieldin purification and proteomic elucidation strategy. Briefly, *Shld1/3* knockout CH12F3 cell-lines were Lentivirally transduced with TwinStrep-HA-Shld1/3-espression transgenes. 300 ml suspension cultures were either mock treated or irradiated (10 Gy) and lysates prepared following a 1hr recovery. shieldin complexes captured from lysates on MagStrep-XT resin were eluted in biotin, captured and tryptic digested on S-Trap columns, with resulting peptides analysed by LC-MS/MS. **B)** B cell SHLD3 vs control (beads-only) interactomes as defined by LC–MS/MS and Label Free Quantification (LFQ). Scatter plot depicts log2 fold-enrichment of indicated MagStrep-XT purified complexes across 2 independent experiments. **C)** Comparison of mock and irradiated B cell SHLD3 interactomes as defined by LC–MS/MS and LFQ. Scatter plot depicts log2 fold-enrichment of indicated MagStrep-XT purified complexes across 2 independent experiments. **D)** Immunoblot analysis of CH12F3 cell extracts from SHLD3 pulldown experiments. Representative of *n = 2* independent experiments.

Highly-efficient purification of shieldin was confirmed by high peptide coverage across all four shieldin proteins (refer to Table 1). Surprisingly, shieldin complexes were notably devoid of significant protein interactors (Fig. 2B). Since interactions between shieldin’s upstream regulators 53BP1 and RIF1 are controlled by DNA damage-dependent phosphorylation^7,8^, our subsequent investigation focused on SHLD1/3 complexes obtained from lysates of X-ray irradiated cells.

**Table 1:** Label-free quantification of LC-MS/MS results. Interacting proteins were determined as in Fig. 2a-c. n=2 independent experiments/genotype.

| Protein Symbol | Unique peptides | Coverage (%) | TS-HA-SHLD1 |  | TS-HA-SHLD3 |  |
| --- | --- | --- | --- | --- | --- | --- |
|  |  |  | LFQ | LFQ<sup>IR</sup> | LFQ | LFQ<sup>IR</sup> |
| Rev7 (MAD2L2) | 8 | 32 | 6.6 | -1.0 | 1.6 | 0.0 |
| Shld1 (C20orf196) | 10 | 41 | 5.1 | 1.2 | 3.1 | -0.3 |
| Shld2 (FAM35A) | 46 | 60 | 3.4 | -0.5 | 5.3 | -0.2 |
| Shld3 (FLJ26957) | 14 | 66 | 2.6 | -0.4 | 4.2 | -0.5 |
| Ctc1 | 1 | - | -3.7 | 8.7 | -0.7 | 3.9 |
| Stn1 | 2 | - | 5.7 | 2.7 | 3.4 | 4.7 |

Remarkably, analysis of (TwinStrep-SHLD1/3) purifications under these conditions revealed a discernible enrichment of peptides from CST proteins CTC1 and STN1 in both sets of purifications (Fig. 2C; Extended Data Fig. 2D). The induction of shieldin-CST interactions following DNA damage treatments was validated through immunoblotting of shieldin complex purifications on a smaller scale using STN1 antibodies (Fig. 2D). Additionally, pre-treatment of cells with a small-molecule ATM inhibitor (KU55933), but not an ATR inhibitor (AZD6738), abolished shieldin-CST interactions (Extended Data Fig. 2E).

These findings strongly implicate shieldin as a recruitment platform for the CST complex during DNA repair and point to ATM-dependent DNA damage signalling as the trigger for assembling shieldin-CST complexes at DSB sites. Such a conclusion provides a likely explanation for the equivalent effects of *shieldin* and *CST* gene disruption in murine B cells, and furthermore implicates Polɑ-Primase-dependent fill-in synthesis as an intermediary step in the joining of AID-induced DSBs.

### Shieldin-dependent DSB repair involves the REV3L polymerase

During DNA replication, the synthesis of an RNA-DNA primer on the lagging-strand by Polɑ-Primase precedes strand extension by DNA Polymerase δ (Polδ)^37^. We therefore considered that processive fill-in synthesis downstream of CST-directed Polɑ-Primase priming might likewise rely on Polδ or an analogous processive DNA polymerase. Interestingly, DNA polymerase ζ (Polζ), a translesion synthesis polymerase formed of the B-family polymerase REV3L in complex with REV7 and Polδ accessory subunits POLD2 and POLD3^38^, was previously implicated in NHEJ during CSR^39^. This led us to investigate REV3L’s participation in DNA end-joining downstream of shieldin/CST.

*Rev3l* is essential in mice^38^, and prior characterisations of B cell-specific REV3L functions were complicated by mosaicism driven by the out-competition of proliferation-defective *Rev3l-*deleted mature B cells with those that had escaped *CD21-Cre*-mediated recombination^39^. We thus opted to breed mice harbouring the same *Rev3l^tm1(Rsky)^* conditional knockout allele in combination with the B cell specific *Mb1-Cre* transgene, which mediates higher-efficiency *Cre-*mediated recombination upstream in B lymphocyte progenitor cells^11,40^. As predicted, *Rev3l* was completely deleted in the B cells of *Rev3l^F/F^ Mb1*^+/Cre^ mice (Extended Data Fig. 3A), where it resulted in 2-3-fold reductions in absolute B cell frequencies in the bone marrow and spleen (Fig. 3B). Despite this, purified B splenocytes from *Rev3l^F/F^ Mb1*^+/Cre^ mice were viable in culture and could undergo multiple rounds of cell division upon stimulation *ex vivo* (Fig. 3C). This allowed us to re-evaluate REV3L’s contribution to CSR. Indeed, *Rev3l^F/F^ Mb1*^+/Cre^ splenic B cells supported IgM to IgG1 class-switching, yet at levels ∼40% lower than wild type or *Mb1-cre-*positive controls (Fig. 3D-F; Extended Data Fig. 3B). Again, class-switching in cell-division staged *Rev3l-*deficient B cells and their controls, confirmed that these reductions were not a consequence of reduced cell proliferation (Fig. 3C and 3D), a result consistent with REV3L’s likely participation in NHEJ^39^.

**Fig. 3:**
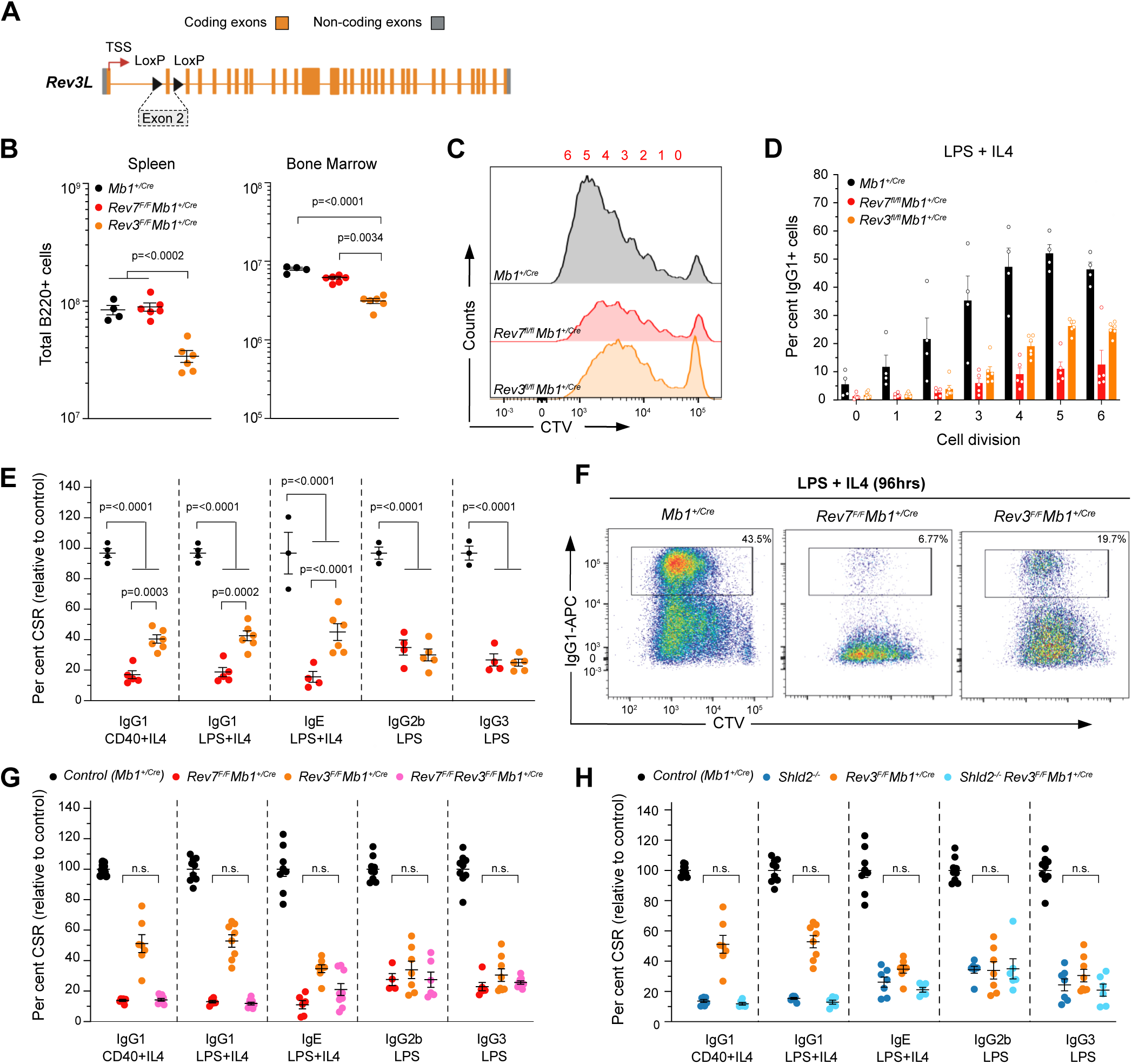
Polζ cooperation with shieldin supports NHEJ in CSR. **A)** Schematic representation of the Rev*3* conditional allele. **B)** Absolute numbers of B220+ B cells in the bone marrow (one femur and one tibia) and spleen. *n = 4-6* mice per genotype. Significance was determined by unpaired two tailed *t*-test, Mean ± SEM. **C)** CTV dilution in purified B cells cultured in the presence of LPS and IL-4 for 96 h. Representative data, *n* > 6 mice. **D)** IgG1-positive (IgG1^+^) B cells as a proportion of total B cells (%) for each cell generation as determined by CTV staining and proliferation-associated dye dilution. *n = 4-6* mice per genotype. Mean ± SEM. **E)** Splenic B cells cultured with the indicated stimuli (96 h) and stained for surface IgG1, IgE, IgG2b or IgG3. *n = 4-7* mice per genotype. CSR 100%, mean immunoglobulin isotype switch frequency of 2 control animals in each experiment. Significance was determined by two-way ANOVA with Tukey’s correction. Mean ± SEM. **F)** Cell trace violet (CTV)-labelled splenic B cells were stimulated as indicated and stained for surface IgG1 after 96 h. Representative data, *n > 6* mice. **G)** Splenic B cells cultured with the indicated stimuli (96 h) and stained for surface IgG1, IgE, IgG2b or IgG3. *n = 4-7* mice per genotype. CSR 100%, mean immunoglobulin isotype switch frequency of 2 control animals in each experiment. **F)** Splenic B cells cultured with the indicated stimuli (96 h) and stained for surface IgG1, IgE, IgG2b or IgG3. *n = 4-7* mice per genotype. CSR 100%, mean immunoglobulin isotype switch frequency of 2 control animals in each experiment.

To ascertain whether Polζ functions downstream of shieldin in CSR, we interbred *Rev3l^F/F^ Mb1*^+/Cre^ mice with *Shld2-*knockout mice, and our previously described B cell conditional *Rev7-*knockout strain^11^, generating mice co-deficient for shieldin and REV3L in the B cell lineage. The *ex vivo* stimulation of mature B splenocytes co-deleted of *Rev3l* and either shieldin gene confirmed that *Rev3l* loss did not further decrease class-switching below that seen in *Rev7 or Shld2* single-knockout controls (Fig. 3G and 3H). This genetic epistasis between *Rev3l* and *shieldin* confirms Polζ’s participation in shieldin/CST-dependent DNA end-joining reactions. Considering CST’s role in catalysing Polɑ-Primase-dependent fill-in synthesis in *BRCA1-*deficient cells and at Cas9-induced DSBs^27,29,30^, our results position Polζ as a likely mediator of processive fill-in synthesis downstream of these DNA priming reactions.

### REV7 and NHEJ -independent REV3L activities supports lymphocyte development

Despite the proliferative capacities of *Rev3l^Δ/Δ^ Mb1*^+/Cre^ stimulated mature B splenocytes, the staging of B lineage cells in the bone marrow of *Rev3l^F/F^ Mb1*^+/Cre^ mice revealed developmental abnormalities. In these mice, diminished frequencies in pro B cell stage cells persisted throughout the small pre-B cell, immature B cell and mature circulating B cell stages, and were noticeably more severe than those observed in *53bp1*^-/-^ mice (Fig. 4A; Extended Data Fig. 4A-C, 1H and 1N). This also contrasted with *Rev7^F/F^ Mb1*^+/Cre^ mice, where Mb1-cre mediated deletion of *Rev7* in B cells caused no development phenotypes (Fig. 4A)^11^, confirming that Polζ’s function in developing B cells requires its polymerase REV3L, but not its accessory subunit REV7. To determine the cause of B cell attrition in *Revl3^F/F^ Mb1*^+/Cre^ mice, we generated *Rev3l^F/F^ p53^F/F^ Mb1*^+/Cre^ mice (Extended Data Fig. 4D). p53 deletion is known to suppress DNA damage-dependent apoptosis in murine lymphocytes^41^, and likewise its disruption rescued B cell frequencies in the bone marrow and spleen of *Rev3l^F/F^ p53^F/F^ Mb1*^+/Cre^ mice (Fig. 4B-C; Extended Data Fig. 4E-G). By contrast, the expression of a transgenic B cell receptor specific for hen egg lysosome (*MD4^+/Tg^*) did not rescue B cell frequencies in *Rev3l^F/F^ Mb1*^+/Cre^ mice, but did so in a V(D)J recombination-defective *53bp1^-/-^* background (Fig. 4D; Extended Data Fig. 4H). Based on this, we surmise that in the *Rev3l-*deleted background B cell lineage abnormalities are most likely precipitated by replication-associated DNA damage, and not by defects in V(D)J recombination. Thus, Polζ supports B lymphocyte development in a REV7 and DNA end joining -independent manner. The rescue of B cell development conferred by p53-deletion is consistent with Polζ’s known role in suppressing DNA replication-associated genomic stability^42^, providing new evidence of REV7-independent functions of Polζ in genome maintenance.

**Fig. 4:**
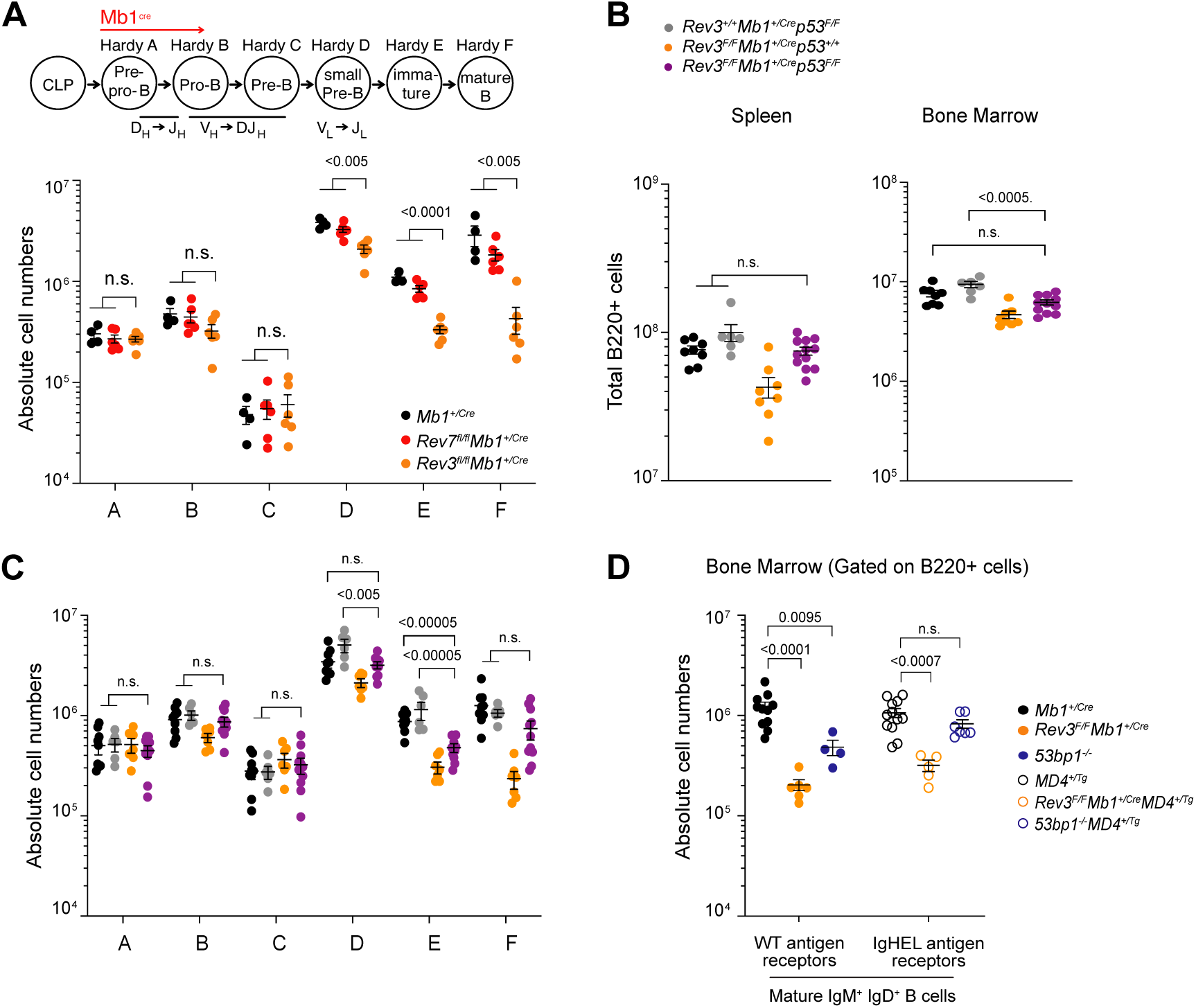
REV7- and NHEJ-independent functions of Polς support B cell development. **A)** Absolute numbers of B cell precursors (Hardy fraction A, B220^+^CD43^+^BP-1^−^CD24^−^; Hardy fraction B, B220^+^CD43^+^BP-1^−^CD24^+^; Hardy fraction C, B220^+^CD43^+^BP-1^+^CD24^+^; Hardy fraction D, B220^+^CD43^−^IgM^−^IgD^−^; Hardy fraction E, B220^+^CD43^−^IgM^+^IgD^−^; and Hardy fraction F, B220^+^CD43^−^IgM^+^IgD^+^) in the bone marrow (one femur and one tibia). *n = 4-6* mice per genotype. Significance was determined by unpaired two tailed *t*-test, Mean±SEM. **B)** Absolute numbers of B220+ B cells in the bone marrow (one femur and one tibia) and spleen. *n = 6-12* mice per genotype. Significance was determined by unpaired two tailed *t*-test, Mean ± SEM. **C)** Absolute numbers of B cell precursors in the bone marrow (one femur and one tibia). *n = 6-12* mice per genotype. Significance was determined by unpaired two tailed *t*-test, Mean ± SEM. **D)** Absolute numbers of Mature IgM^+^ IgD^+^ B cells in the spleen. n ≥ 4 mice per genotype, Mean ± SEM.

### Minimal contribution of shieldin to developmental defects in *BRCA1-*knockout mice

The above findings confirm the 53BP1 pathway directs multiple, mechanistically-distinct activities, which can be distinguished by shieldin-CST involvement. Synergy between these activities is required for the efficient joining of AID-induced DSBs during CSR, yet during V(D)J recombination 53BP1 acts independently of shieldin, CST or REV3L. We reasoned that these activities might therefore each contribute separately to the pathological events in *Brca1-*deficient mice and tumours, where the 53BP1 pathway impedes HR, and through shieldin and CST, mediates toxic NHEJ events^11,16,21,22,24,26,27,29^. To test this, we interbred *53bp1* and *Shld2* knockout mice with mice harbouring *Brca1* exon 5-13 deletions (*Brca1^+/-^*) that ablate all BRCA1 protein expression^43^. *Brca1^-/-^* mouse embryos die early in development, with none outlasting embryonic day E7.5^44^. However, embryonic lethality can be circumvented upon deletion of *53bp1*, yielding viable, albeit tumour-prone mice^19,20^. In our hands, *Brca1^-/-^ 53bp1^-/-^* mice were born at around half of the expected frequency (Fig. 5A and Table 2), and weighed less than their *Brca1^+/-^53bp1*^-/*-*^ littermates, with most succumbing to thymic lymphoma between 12-20 weeks of age (data not shown), consistent with recent reports^20,45^. By contrast, no *Brca1^-/-^ Shld2^-/-^* double-knockout pups were born to intercrosses between *Brca1^+/-^ Shld2^-/-^* mice (Table 2). Furthermore, we recovered only non-viable *Brca1^-/-^Shld2*^-/-^ embryos between stages E8.5-E11.5, all of which exhibited severe developmental delay (Fig. 5B and Table 2). Of note, from staged embryos obtained in *Brca1* heterozygous intercrosses performed in parallel (analysed at E8.5 and E10.5), no *Brca1^-/-^* embryos were recovered. This implies that unlike the case for *53bp1-*deficiency, *shieldin*-inactivation cannot restore viability in *Brca1* null mice, and only modestly delays the onset of embryonic lethality.

**Fig. 5:**
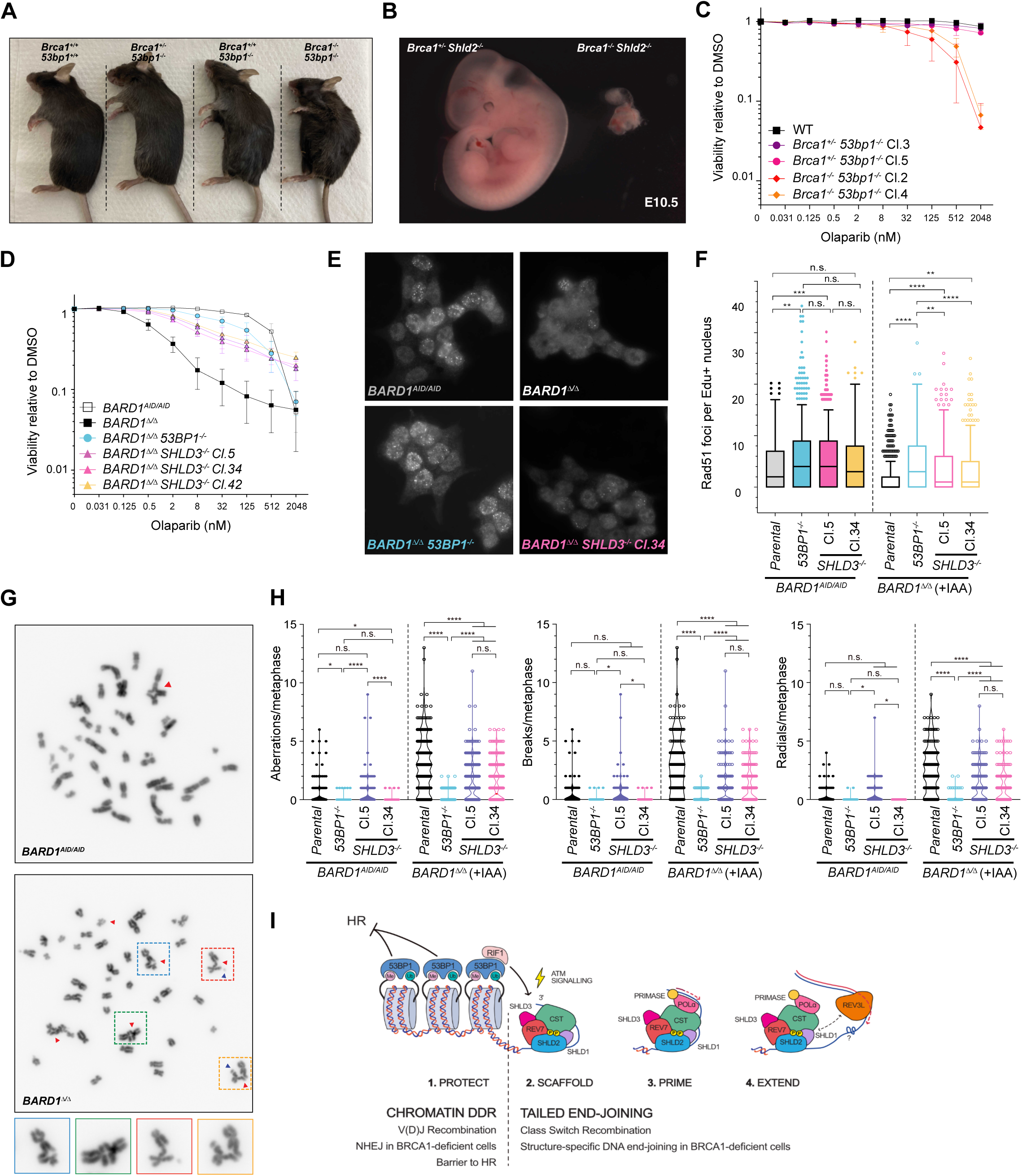
Independent and synergistic contribution of 53BP1 and shieldin to HR suppression and NHEJ in *BRCA1-*deficient cells. A) Representative image of *53bp1^-/-^Brca1^−/−^* mice and relevant controls at ∼12 weeks. Genotypes as indicated. B) Representative image of *a Shld2^-/-^ Brca1^+/−^* control embryo and *Shld2^-/-^ Brca1^−/−^* double KO embryo at E10.5. C) Survival of the indicated MEF cell lines grown for 7 days in the presence of the indicated doses of Olaparib, *n=3* biological experiments, Mean±SD. D) Survival of the indicated *BARD1^AID/AID^* HCT-116 cell-lines grown for 7 days in the presence or absence of IAA (1mM), doxycycline (2 μg ml^-1^) and the indicated doses of Olaparib, *n=3* biological experiments, Mean±SD. E) Immunofluorescent microscopy of Rad51 IRIF in Edu^+^ indicated BARD1^AID/AID^ cell lines. Cultures were supplemented with doxycycline (2 μg ml^-1^ for 24 h) before addition of IAA (1 mM). After 24h cells were irradiated (5 Gy) and fixed 2h later. Representative images of n=3 biological experiments. F) Quantification of RAD51 IRIF from E. Integrated intensity and foci quantifications were made using CellProfiler. Boxes indicate the 25th–75th percentiles with the median denoted, and whiskers calculated by the Turkey method. Significance was determined by two-sided Kruskal–Wallis H test with Dunn’s correction for multiple comparisons. ****P<0.0001, **P <0.002. G) Representative metaphase images from the indicated *BARD1^AID/AID^* cell lines. Cultures were supplemented with doxycycline (2 μg ml^-1^for 24h) before addition of IAA (1 mM). After a further 24h cells were treated with 500nM Olaparib and fixed 24 h later. Representative images of *n=3* biological experiments. H) Quantification of between 40 and 65 metaphases analysed per genotype/experiment. Significance was determined by two-sided Kruskal–Wallis *H* test with Dunn’s correction for multiple comparisons. *\*\*\*\*P<0.0001, ***P<0.0002, **P<0.002, *P<0.05*. I) Model for 53BP1-dependent DSB repair. Chromatin associated 53BP1 complexes suppress HR and promote V(D)J recombination. DNA damage dependent signalling via RIF1 is conferred to shieldin via interactions with SHLD3. DSB signalling via ATM stimulates the assembly of shieldin-CST complexes on ssDNA, promoting the recruitment of Polɑ-Primase to initiate fill-in synthesis. Polζ polymerase REV3L increases the processivity of Polɑ-Primase catalysed fill-in synthesis.

**Table 2:** *Brca1^-/-^* intercrosses. Summary of the frequency of live pups born or >E12.5 embryos genotyped from *53bp1*^-/-^ *Brca1*^+/−^ or *Shld2*^-/-^*Brca1*^+/−^ crosses.

| *Brca1<sup>+/-</sup> 53bp1<sup>-/-</sup> x Brca1<sup>+/-</sup> 53bp1<sup>-/-</sup>* |  |  |  |
| --- | --- | --- | --- |
| *Brca1* | +/+ | +/- | -/- |
| *53bp1* | +/+ | -/- | -/- |
| Live pups<br>(n=209) | 68<br>(52.25) | 120<br>(104.5) | 21<br>(52.25) |
| *Brca1<sup>+/-</sup> Shld2<sup>-/-</sup> x Brca1<sup>+/-</sup> Shld2<sup>-/-</sup>* |  |  |  |
| *Brca1* | +/+ | +/- | -/- |
| *Shld2* | +/+ | -/- | -/- |
| Live pups<br>(n=242) | 89<br>(60.5) | 153<br>(121) | 0<br>(60.5) |
| Embryos<br>(E8.5-11.5)<br>(n=57) | 15<br>(14.25) | 32<br>(28.5) | 10<br>(14.25) |

### 53BP1 controls DSB repair pathway choice independently of shieldin

As shieldin loss blocked CSR yet could not rescue the development of *Brca1-*null embryos, we hypothesised that the pro-NHEJ and HR-suppressive activities of the 53BP1 pathway might be distinguished at the level of shieldin-involvement. Suggestive of 53BP1 operating independently of shieldin during HR-suppression, we were unable to derive viable embryonic fibroblast (MEF) or embryonic stem cell (mESC) cell-lines from *Brca1^-/-^ Shld2*^-/-^ E8.5 embryos and blastocyst outgrowths, respectively. We therefore resorted to addressing shieldin’s involvement in engineered *BARD1^AID/AID^* HCT-116 cells where BRCA1 can be conditionally inactivated through auxin degron-mediated degradation of its obligate binding partner protein BARD1^14,15^. Direct binding of DSB-proximal nucleosomes ubiquitinated on H2A Lysine 15 (H2A K15ub) by BARD1 is a pre-requisite for BRCA1-dependent pathway choice control, since it directly inhibits 53BP1-chromatin interactions at DNA damage sites^14,17^. Accordingly, in auxin-treated *BARD1^AID/AID^* HCT-116 cells (here forth referred to as *BARD1^τ./τ.^* cells), HR is fully restored upon deletion of 53BP1, resulting in a dramatic (albeit incomplete) suppression of cellular hypersensitivity to PARPi Olaparib^14^. Likewise, mouse embryonic fibroblast cell-lines derived from *Brca1^-/-^53bp1^-/-^* embryos exhibited similar levels of residual Olaparib sensitivity (Fig. 5C and 5D), reassuring us of the suitability of *BARD1^AID/AID^* HCT-116 as a cellular model to investigate and compare the contributions of 53BP1 and shieldin to HR defects.

In line with previous results in *BRCA1-*deficient cells^11,22,26^, *SHLD3-*deletion in *BARD1^τι/τι^* cells markedly enhanced cell survival in Olaparib cytotoxicity assays. Importantly, however, olaparib resistance was less penetrant in *BARD1^τι/τι^ SHLD3^-/-^* cells, as evidenced by the enhanced relative survival of *BARD1^τι/τι^ 53BP1^-/-^* cells in these experiments (Fig. 5D; Extended Data Fig. 5A). HR defects in replicating (S phase staged) *BARD1^τι/τι^ SHLD3^-/-^* cells were also evident at the level of defective RAD51 recruitment into IR-induced foci, in stark contrast to *BARD1^τι/τι^ 53BP1^-/-^* cells, where RAD51 IRIF accumulated at wild type (non-auxin treated *BARD1^AID/AID^* cells) levels (Fig. 5E-F).

The different extents to which inactivation of 53BP1 or shieldin could suppress DNA repair defects in BRCA1-deficient (*BARD1^τι/τι^*) cells was also evident at the level of chromosome stability. In metaphase chromosome spreads from *BARD1*^τι/τι^ cells, olaparib treatment induced high frequencies of chromosome/chromatid breaks and radial chromosomes (Fig. 5G-H; Extended Data Fig. 5B). Chromosome breaks and radials were only slightly reduced in frequency in *BARD1*^τι/τι^ *SHLD3*^-/-^ cells, highlighting 53BP1’s retained capacity to block repair, and mediate toxic joining events independently of shieldin. By contrast, both classes of chromosome lesion were near-completely suppressed in *BARD1*^τι/τι^ *53BP1*^-/-^ cells (Fig. 5G-H), consistent with their restored capacity for HR (Fig. 5E-F). We can therefore conclude that HR inactivation in *BRCA1/BARD1-*deficient cells is a consequence of 53BP1, but not of shieldin or its cooperating proteins. Nevertheless, in the absence of BRCA1-dependent HR promotion, shieldin-directed DNA end joining exacerbates DNA damage-induced chromosomal instability, where it also makes substantial contributions to the overall hypersensitivity of cancer cells to HRD-targeting PARPi.

## DISCUSSION

In this study, we adopted a genetic approach to systematically dissect the 53BP1 pathway. By comparing the traits of mice deficient in 53BP1, shieldin, or CST, we aimed to elucidate the intricate relationship between 53BP1 and its downstream DNA processing components in both normal and pathological contexts. Our findings shed light on the remarkable capability of 53BP1 to facilitate DSB repair even in the absence of shieldin. However, they also establish the indispensability of shieldin-CST interplay for efficient 53BP1-dependent DNA end-joining during CSR, and furthermore confirms Pol(‘s involvement in these important DNA synthesis-dependent DNA end-joining reactions.

### Shieldin is a DSB repair-specific mediator of CST

Upon binding ssDNA at telomeres and during DNA repair, CST acts as a catalyst of Polɑ-Primase-dependent DNA synthesis^29,46^. The discovery of shieldin-CST interdependence during CSR thus underscores the specialized nature of fill-in DNA synthesis as a prerequisite for the joining of AID-dependent DSBs. We additionally demonstrate that 53BP1’s reliance on shieldin/CST differs dramatically across different repair contexts. We posit that these differences reflect the unique DNA end-processing demands inherent to the spectrum of DSB structures generated in each distinct scenario, coupled with their varying frequencies of occurrence.

These conclusions are supported by multiple parallel observations in lymphocytes, mice, and cancer cell models. Besides the equivalent severity of class-switching defects in mature B cells from *Shld2^-/-^, Shld3^-/-^* and *Ctc1^F/F^ Mb1^+/cre^* mice, hypothesis-agnostic elucidation of the B cell shieldin-interactome in B cells identified CST proteins as its primary interaction partners. Interestingly, interactions between shieldin and CST are induced upon DNA damage, a behaviour explained by its reliance on the DSB-responsive kinase ATM. Upstream of shieldin, ATM-dependent phosphorylation of consensus Ser-Glu containing motifs in 53BP1 promote the recruitment of RIF1^7,47^ via a recently discovered phospho-ligand binding module in the RIF1 heat repeats^8^. Given the absence of additional shieldin interactors in our proteomics datasets, the ATM-dependent phosphorylation events that guide shieldin-dependent CST recruitment to DNA damage sites most-likely involves a cryptic phospho-ligand binding module within either shieldin or CST. From a structural perspective, it is noteworthy that neither CSR, nor the ATM-dependent shieldin-CST interaction discovered here, relies on a recently described N-terminal CTC1-binding motif in SHLD1^29^. Indeed, we found that a SHLD1^Δexon1^ protein deleted of its first 59 amino acids (encompassing this motif at a.a. 18-21) efficiently retrieved CST from lysates when they were prepared from irradiated B cells (Extended Data Fig. 2F). Similarly, mature splenic B cells from *Shld1^Δexon1^* mice (a gift from the MMRRC^48^) confirmed to express an analogous truncated protein were also found to support near-wild type class-switching frequencies in *ex vivo* stimulation experiments (Extended Data Fig. 2G-H), consistent with our previous results in *Shld1^-/-^* CH12F3 cells expressing a *Shld1^ΔN^* transgene^29^. Nevertheless, ATM-dependent shieldin-CST interactions are similarly consistent with shieldin’s importance in recruiting CST to DSB termini, explaining their inter-dependence in downstream DNA end-joining.

Interestingly, this concept draws intriguing parallels with the regulation of CST-dependent C-strand fill-in synthesis at telomeres, governed by the shelterin complex. In the telomeric context, the recruitment of CST to telomeres hinges on direct interactions between CST and the shelterin protein POT1b^46^. Recent structural findings reveal this involves a multi-site interaction between CTC1 and regions within the triple OB-fold domain in POT1, which is further potentiated by POT1 phosphorylation^49^. A similar triple OB-fold architecture is also present in SHLD2^11,21,22,26^, and so it is tempting to speculate that a conserved mode of binding between CTC1 and structurally-related features in POT1 or SHLD2 might underpin the initiation of fill-in DNA synthesis at telomeres, and DSBs, respectively^27,29,46^.

### Polζ acts downstream of shieldin-CST during tailed-end joining

Using mice co-deleted for *Rev3l* and either *Shld2* or *Rev7,* we additionally detected an epistatic relationship between these genes in class-switching mature B cells. This offers a rational mechanistic explanation for Pol(’s role in NHEJ^39^, positioning it downstream of shieldin/CST, and most likely in the processive DNA repair synthesis that follows priming by Polɑ-Primase. Class-switching in *Rev3l-*deficient B cells occurs at ∼50% of wild type frequencies, a level ∼3-4 fold higher than that seen in those lacking *shieldin* genes or *Ctc1*. Does this indicate that Polζ is not the sole DNA polymerase acting downstream of CST-Polɑ-Primase? As the typical extender of lagging-strand primers during DNA replication, Polδ would likely be a redundant polymerase. It is also possible that RNA-DNA primer synthesis on a short 5′-resected DNA end might be sufficient to stimulate the ligation of ssDNA-tailed DSBs by NHEJ, a pathway able to ligate DSBs with ribonucleotide-incorporated termini^50^. Nevertheless, AID’s action during CSR generates ssDNA-tailed DSBs within switch *(S)* regions in the *IgH* locus, GC-rich repeat-dense intronic elements abundant in palindromic motifs. The propensity of *S*-region ssDNA to form stem loops and/or parallel four-stranded G-quartets is well-documented, and such properties are speculated to contribute to various aspects of *S-*region function during CSR^51^. The formation of secondary DNA structures in the ssDNA-tailed termini of AID-dependent DSBs might similarly impede DNA synthesis by the canonical lagging-strand polymerase Polδ, warranting the utility of a versatile TLS polymerase to mediate DNA fill-in synthesis at *IgH* DSBs. Polζ also limits DNA damage accumulation during normal replication^42^, a function we suggest extends to developing B cells that surprisingly doesn’t involve REV7. Whether Polζ participates in other CST-Polɑ-Primase catalysed fill-in reactions remains to be explored. Such an interplay might be advantageous during C-strand fill-in at telomeres, where secondary DNA-structures on the template G-strand could pose similar challenges to Polδ processivity^52^.

### 53BP1 promotes DNA end-joining independently of shieldin-CST

Based on the genetic analyses described above, we propose a refined model for the shieldin-dependent branch of the 53BP1 pathway (Fig. 5I). While vital for optimal CSR, the residual switching capacity of stimulated shieldin/CST-deficient B splenocytes significantly exceeds that in *53bp1*-deficient B cells, confirming 53BP1 promotes the repair of a small subset of AID breaks without the requirement for fill-in synthesis. An additional paradigm for shieldin-independent NHEJ is V(D)J recombination, where 53BP1 supports the development of normal frequencies of B and T lymphocytes by enhancing long-range joining events^10,11,13^. Why does 53BP1 operate independently of shieldin/CST during V(D)J recombination? The simple explanation for this is RAG-induced DNA breaks have no utility for shieldin/CST-dependent fill-in synthesis yet profit from a distinct activity exerted by 53BP1. Our comparative investigations of 53BP1 and shieldin functions in *BRCA1*-deficient cells provide additional evidence in support of this proposition: shieldin-inactivation confers a degree of PARPi resistance to *BARD1-*depleted cells, yet resistance was further increased by deletion of 53BP1. Similar distinctions were perhaps hinted at by murine tumour studies: the potent survival benefit conferred by PARPi treatments in mice implanted with *Brca1^-/-^ Trp53^-/-^* mammary tumours was completely lost when tumours were additionally deleted of *53bp1*^53^; however, they modestly extended lifespan in equivalent tumour engraftment experiments when the tumours were instead deleted for *Shld1* or *Shld2*^22^. Our knockout experiments in *BARD1^Δ/Δ^* cells also reveal 53BP1 to be the predominant mediator of PARPi-induced chromosome breaks and radial chromosomes, with shieldin making a lesser contribution. Given the different contributions that 53BP1 and shieldin make to the repair of RAG- and AID-induced DSBs in lymphocytes, we speculate that DSBs resulting from PARPi-induced replication blocks might likewise comprise a spectrum of DNA-end structures, of which only some will necessitate shieldin-CST axis-dependent repair.

### HR suppression is an upstream, shieldin-independent, function of the 53BP1 pathway

Among the most striking deviations in phenotypes conferred by 53BP1 and shieldin losses, was at the level of HR modulation in *BRCA1-*deficient (*BARD1^Δ/Δ^*) cells. Here, 53BP1 deletion fully restored RAD51 recruitment to wild-type levels, indicative of fully-competent HR. By contrast, profound RAD51 recruitment defects in *BARD1^Δ/Δ^* cells similarly presented in *shieldin-*inactivated derivative cell-lines, a distinction that implies that DSB repair pathway choice regulation by 53BP1 is agnostic to shieldin/CST mediated DNA end-processing. The weak degree of observed phenotypic suppression in *Shld2^-/-^ Brca1^-/-^* embryos offers the most compelling support for this conclusion: shieldin-disruption only slightly delayed the onset of embryonic lethality, in stark contrast to the viability and superficially normal post-natal development of *53bp1^-/-^ Brca1^-/-^* mice. These findings may seem in contradiction to a recent report, where *Shld2* knockout rescued embryonic and post-natal development in mice homozygous for the hypomorphic *Brca1^Δ^*^11^ mutation^12^. However, these differences are almost certainly attributable to the significant residual functionality of the BRCA1^Δ^^11^ protein. BRCA1^Δ11^-BARD1 complexes are readily recruited to sites of DNA damage where they support functional interactions with HR mediators such as PALB2-BRCA2. However, DNA end-resection is severely reduced in cells deleted of exon 11 in *BRCA1*^54^, representing a cellular defect that might interfere with HR when challenged with antagonistic 53BP1-directed shieldin/CST fill-in activities.

In summary, our study reveals a surprising segregation-of-duties between 53BP1 and its downstream effector proteins during DSB repair. Shieldin/CST-dependent fill-in synthesis at DNA ends constitutes an auxiliary reaction during the joining of resected DNA ends, such as those generated at AID-dependent DSBs during CSR. However, outside of CSR, 53BP1-dependent DSB repair remains remarkably resilient to loss of these effectors, consistent with a direct stimulation of DNA repair by chromatin-bound 53BP1 complexes. Our results therefore implicate upstream 53BP1 complexes, but not its DNA processing effectors, as the major determinants of HR suppression in mammalian cells (Fig. 6I). This concept aligns well with our recently proposed chromatin-centric model for DSB pathway choice control, where HR-NHEJ outcomes are dictated by a mutual antagonism between BRCA1-BARD1 and 53BP1, upon their engagement of related DSB-proximal chromatin signatures^14,15^. Future work will be needed to address whether this hierarchy in the 53BP1 pathway will also be reflected in the clinical penetrance of mutations affecting *53BP1* and its effector genes in drug-resistant *BRCA1* mutant cancers.

## Data and code availability

The mass spectrometry proteomics data have been deposited to the ProteomeXchange Consortium via the PRIDE [1] partner repository with the dataset identifier PXD045534 and 10.6019/PXD045534.

## Author contributions

Conceptualization, A.K., J.R.C.; Methodology, A.K., R.P., C.O., J.R.B., D.B., C.P., B.D., J.R.C.; Investigation, A.K., P.R., J.S.M., R.P., D.M., C.O., B.D.; Formal Analysis, A.K., R.P.; Data Curation, R.P.; Writing – original draft, review & editing, A.K., J.R.C.; Funding Acquisition, J.R.C; Supervision, A.K., J.R.C.

## Acknowledgements

We thank all members of the laboratory of J.R.C. for discussions; Jos Jonkers for *Brca1/p53* mouse models; Shan Zha, Barry Sleckman & Bo-Ruei Chen for advice, reagents and cell-lines, Mukta Deobagkar for technical assistance; the European Mouse Mutant Cell Repository (www.eummcr.info) and Mutant Mouse Resource & Research Centers (www.mmrrc.org) for mouse strains and gene-targeting constructs. University of Oxford Department of Biomedical Services (BMS) Functional Genetics and JR Hospital BMS facilities for technical support; This work was funded by Medical Research Council (MRC, UK) project grant MR/R017549/1 and Medical Research Council (MRC, UK) Molecular Haematology Unit grants MC_UU_00016/19 and MC_UU_00029/2; Salary to J.RC. and funding for his group comes from a Cancer Research UK (CRUK) Senior Cancer Research Fellowship (RCCSCF-Nov21\100004) and previously, Career Development Fellowship (C52690/A19270) which provided salary support to C.O. and J.R.B.; A.K. and J.R.B. were previously supported by EMBO Long-Term (EMBO - ALT 542-2020) and Ruth L. Kirschstein NRSA Individual Postdoctoral Fellowships (F32) (NIH/NCI - F32CA239339), respectively. The international Mouse Phenotyping Consortium (IMPC) supported this work via UK Research Institute grant MC_UP_1502/3. This work was previously supported by the Wellcome Trust (Core Award Grant, 203141/Z/16/Z). J.R.C. is a recipient of a Lister Institute Research Prize Fellowship.

## Methods

### Mice

The production and breeding of the genetically altered mice was carried out in accordance with UK Home Office Animal (Scientific Procedures) Act 1986, with procedures reviewed by the Clinical Medicine Animal Welfare and Ethical Review Body at the University of Oxford, and conducted under project license PP8064604. Animals were housed in individually ventilated cages, provided with food and water ad libitum and maintained on a 12 h light:12 h dark cycle (150–200 lx). The only reported positives on FELASA health screening over the entire time course of these studies were for *Helicobacter hepaticus and Entamoeba* spp. Experimental groups were determined by genotype and were therefore not randomized, with no animals excluded from the analysis. Sample sizes for fertility studies were selected on the basis of previously published studies and all phenotypic characterization was performed blind to experimental group. All mice used in this study were generated on or backcrossed onto a C57BL/6 Background (> 5 generations). *Shld2^Tm1a^* (*Shld2^tm1a(EUCOMM)Hmgu^*, MGI:5428631) mice generated by the European Conditional Mouse Mutagenesis Program for the International Mouse Knockout Consortium, were crossed with Flp-Cre positive mice (Tg(*ACTB*-*Flpe*)9205Dym; Jax stock 005703) to generate a conditional allele with LoxP sites flanking exons 4 and 5 which includes the translation start site^35^. A targeted *Shld2*-null allele was subsequently generated by germline recombination of the LoxP sites using PGK-cre^55^ (B6.C-Tg(Pgk1-cre)1Lni/CrsJ; Jax stock 020811) (Fig. 1a). *Shld3*-null mice were generated at MRC Harwell by microinjection of Cas9 ribonucleoprotein complexes targeting exon 2 in the C57BL/6 background. Two founder mice, *Shld3^DEL2061-EM1^* and *Shld3^DEL2061-EM2^*, with deletions of the sole coding exon were generated (2054bp and 2061bp respectively) (Fig. 1a). Conditional *Ctc1^F/F^* mice were generated by crossing the ‘knockout-first’ allele (*Ctc1^tm1a(KOMP)Wtsi/Ieg^;* MGI:4363331) to Flp-Cre positive mice (Tg(*ACTB*-*Flpe*)9205Dym; Jax stock 005703) to generate mice with the *Ctc1^tm1c^* conditional allele. Experimental *Ctc1^F/F^ Mb1*^+/Cre^, *p53^F/F^ Mb1*^+/Cre^ (Trp53^floxed^; MGI:98834) and *Rev3L^F/F^ Mb1*^+/Cre^ (Rev3l^tm1Rsky^; MGI:1337131) mice were generated by intercrossing with mice expressing the B cell lineage *Mb1*-cre deleter strain (*Cd79a^tm1(cre)Reth^;* MGI:368745^40^). *53bp1*^−/−^ mice (*Trp53bp1^tm1Jc^*; MGI:2654201) and *Brca1^−/−^* mice (Brca1^null^; MGI:104537) were generated and described elsewhere^19,43,56^. *Shld1^Δexon1^* mice (*C57BL/6NCrl-1110034G24Rik^em1(IMPC)Mbp/Mmucd^;* stock 043862-UCD), obtained from the Mouse Biology Program (University of California Davis, CA, USA) harboured a CRISPR/Cas9-generated deletion of the terminal 71 bp of *Shld1* exon 1. cDNA sequencing confirmed the expression of a functional spliced *Shld1^Δexon1^* transcript, initiated at an indel-generated in-frame ATG codon within the preserved frame-shifted sequences in exon 1 (ATG encoded within Tyr-29-Glu-30 codons).

All B-cell phenotyping experiments involved age-matched 8–16-week-old male or female animals on an inbred C57BL/6 background. We were not pursuing lower penetrance phenotypes, thus statistically significant data could typically be obtained with small group sizes (typically, 4–10 mice). Randomization of samples was only undertaken during the scoring of chromosomal aberrations during metaphase analyses. Mice of a certain genotype were selected based on a unique mouse ID number that does not indicate mouse genotype, thus phenotype–genotype relationships were determined only at the data analysis stage. All experiments were approved by the University of Oxford Ethical Review Committee and performed under a UK Home Office Licence in compliance with animal use guidelines and ethical regulations.

### Immunisations

Wild-type^+/+^, *53bp1*^-/-^, *Shld2*^-/-^ or *Shld3*^-/-^mice were immunised intraperitoneally with 50 mg of NP-CGG (Santa Cruz Biotechnologies) emulsified in Imject Alum adjuvant (Pierce, Thermo Fisher Scientific). Blood samples were collected from the tail vein at 0, 7, 14, 21 and 28 days after immunization.

### Enzyme-linked Immunosorbent assays

Enzyme-linked immunosorbent assays (ELISAs) were used to quantify the production of NP-specific antibodies in mice serum. Ninety-six-well plates were coated with 1 μg/ml NP-BSA (Biosearch Technologies) in bicarbonate buffer, blocked with 5% milk in PBS and incubated with serial dilutions of serum collected at different time points from immunized mice. Plates were probed using alkaline phosphatase-coupled antibodies against mouse IgM and IgG1 (Southern Biotech). Phosphatase substrate (Sigma) was used for detection and optical density measured at 405 nm. For IgG1, pooled blood from post-immunization wild-type mice was used as a standard and serially diluted into a standard curve. The first dilution was established as 1,000 arbitrary units. For IgM pooled blood from day 7 was used as a standard. Immunoglobulin concentrations in mouse serum or culture supernatants were determined by sandwich ELISA. Total IgG, IgM and IgA was measured with mouse IgG, IgM and IgA ELISA kits, respectively (Bethyl Laboratories), according to the manufacturer’s instructions. Mouse serum with known immunoglobulin concentrations of each immunoglobulin was used as a standard.

### Micronuclei analysis

Micronuclei from peripheral blood samples were analysed as described Balmus et al. Nature Protocols, 2014 (<u>10.1038/nprot.2015.010</u>). ∼50ul of tail-vein blood was collected into heparin solution and immediately fixed in cold methanol. Samples were stored at -80°C until processing.

### Lymphocyte analysis and flow cytometry

Cell suspensions from bone marrow (one femur and one tibia) and spleen were counted on a Pentra ES60 Haematology Analyser (Horiba) and stained with anti-mouse antibodies against the following antigens as appropriate (all from BioLegend unless otherwise stated) in FACS buffer (PBS with 2% BSA and 0.025% sodium azide): IgD (1:500, 405716 or Thermo Fisher 12-5993-82; 11-26c.2a), IgG1 (1:200, 406606; RMG1-1), IgM (1:500, 406506 or 406514; RMM-1), B220 (1:500, 103232, 103244, 103206, or 103212; RA3-6B2), BP-1 (1:200, 108308; 6C3), CD19 (1:500, 115534 or 115520; 6D5), CD24 (1:1000, 101827 or BD Pharmingen 562563; M1/69), CD93 (1:200, 136510; AA4.1), CD23 (1:200, BD Pharmingen 553139; B3B4), CD43 (1:200, BD Pharmingen 562865; S7, or Thermo Fisher 11-0431-85; eBioR2/60), and CD21 (1:500, BD Biosciences 563176; 7G6). Mouse BD Fc Block (1:500, BD Pharmingen 553141) was added to block non-specific binding and live/dead cells were discriminated after staining with Zombie Aqua viability dye (1:200, 423102). Data were acquired on a FASCCanto (BD Biosciemces), LSR Fortessa X20(BD Biosciences) or Attune NxT Flow Cytometer (Thermo Fischer) and were analysed with FlowJo Software v10 (Tree Star). Gating strategies for bone marrow Hardy Fractions are shown in Supplementary Fig. 1 and 5.

### Ex vivo B splenocyte culture, stimulation and flow cytometry

B cells were purified from red blood cell-lysed single-cell suspensions of mouse spleens by magnetic negative selection using a B Cell Isolation Kit (Miltenyi Biotec, 130-090-862). B cells (3x10^5^ per well in a 12-well plate) were cultured in RPMI supplemented with 10% FCS, 100 U/ml penicillin, 100 ng/ml streptomycin, 2 mM l-glutamine, 1× NEM nonessential amino acids, 1 mM sodium pyruvate and 50 μM β-mercaptoethanol. B cells were stimulated with 5 μg/ml LPS (Sigma, L7770-1MG), 10 ng/ml mouse recombinant IL-4 (Peprotech, 214-14-20), and agonist anti-CD40 antibody (0.5 μg/ml; Miltenyl Biotec; FGK45.5). Cultures were grown at 37 °C with 5% CO_2_ under ambient oxygen conditions. Four days after seeding, stimulated B cells were analysed using a FACS Canto or LSR Fortessa X20; analysis was performed using FlowJo. Cells were resuspended in FACS buffer, blocked with Mouse BD Fc Block, and immunostained with biotinylated antibodies as follows: anti-mouse IgG1 (1:100, BD Pharmingen 553441; A85-1), anti-mouse IgG2b (1:100, BioLegend 406704; RMG2b-1), anti-mouse IgG3 (1:100, BD Pharmingen 553401; R40-82) and Streptavidin APC (1:500, Thermo Fisher 17-4317-82). Cells expressing IgE were assessed using anti-Mouse IgE PE (1:200, BioLegend, 406908; RME-1). Live/ dead cells were discriminated after staining with Zombie Aqua viability dye. Cell proliferation was assessed using Cell Trace Violet according to manufacturer’s instructions (CellTrace, Life Technologies).

### Proteomics and Mass spectrometry

TwinStrep-HA-SHLD1/3 or control complexes were isolated from lysates of H12-F3 cells on MagStrep “type3” StrepTactinXT beads washed extensively in wash buffer and eluted with Biotin. TwinStrep-peptide eluted complexes were solubilised in 4% (w/v) SDS, 50 mM TEAB, reduced and alkylated using DTT and iodoacetamide and digested with trypsin on S-Trap micro columns as per the manufacturer’s instructions. Samples were analysed on a LC-MS platform (Ultimate 3000 nHPLC and Q-Exactive HFX mass spectrometer). Peptides were separated on an 50 cm EASY-Spray column with a gradient of 3–35% acetonitrile in 5% DMSO and 0.1% formic acid at 250 nl/min. MS1 spectra were acquired at a resolution of 60,000 at 200 m/z. Up to the 15 most abundant precursor ions were selected for subsequent MS/MS analysis after ion isolation with a mass window of 1.6 Th. Peptides were fragmented by HCD with 27% collision energy. Raw files were analysed by Progenesis QI (v.3, Waters) for spectral Label Free Quantitation with default parameters (top 3 quantitation mode). Proteins were identified with PEAKS 8.0 (Bioinformatics Solutions) using standard parameters and the Uniprot mouse reviewed proteome (retrieved 23 January 2020). Peptide and Protein false discovery rate(s) (FDR) was set to 1%.

### Cell lines, cell culture and CRISPR-Cas9 editing

All CH12-F3 cell lines were cultured in RPMI supplemented with 5% NCTC-109 medium, 10% FCS, 100U/ml penicillin, 100ng/ml streptomycin and 2mM l-glutamine at 37°C with 5% CO2 under ambient oxygen conditions. *Rev7*^-/-^, *c20orf196*^-/-^ (*Shld1*^-/-^) and *Flj26957*^-/-^ (*Shld3*^-/-^) CH12-F3 were generated using CRISPR–Cas9 as previously described^11^. Complemented cell lines were generated by lentivirus-mediated transduction, using viral supernatants collected from 293T cells co-transfected with third generation packaging vectors and a pLenti-PGK-TwinStrep-Flag-HA-PURO-DEST vector containing cloned transgene inserts. Typically, cells were spinoculated with polybrene (8 μg/ml) and HEPES (20 mM)-supplemented viral supernatants (1500 rpm, 90 min at 25 °C). Stable cell-lines were subsequently selected and maintained in the presence of puromycin (1μg/ml). To stimulate CSR to IgA, CH12-F3 cells were stimulated with agonist anti-CD40 antibody (0.5μg/ml; Miltenyi Biotec; FGK45.5), mouse IL-4 (5 ng/μl; R&D Systems) and TGFβ1 (2.5ng/μl; R&D Systems). Cell-surface IgA expression was determined by flow cytometric staining with anti-mouse IgA-PE antibody (Cambridge Bioscience; 1040-09).

E12.5 MEFs prepared and SV40 LargeT-immortalized by standard procedures were used in all experiments unless otherwise indicated. All MEF cell lines were maintained in Dulbecco’s modified Eagle medium (DMEM)–high glucose (Sigma-Aldrich, D6546) supplemented with 10% FBS, penicillin–streptomycin and 2mM L-glutamine.

*BARD1^AID^*^/AID^ cell lines and *BARD1*^AID/*AID*^*53BP1^-/-^* were generated as previously described^14,15^. All *BARD*1^AID/AID^ and derivative cell lines were maintained in Dulbecco’s modified Eagle medium (DMEM)–high glucose (Sigma-Aldrich, D6546) supplemented with 10% FBS, penicillin–streptomycin and 2mM L-glutamine. Cultures were maintained at 37 °C with 5% CO_2_. *BARD1*^AID/*AID*^*SHLD3*^-/-^ knockout cell lines were generated by CRISPR/Cas9 as previously described^14^. Gene-specific gRNAs were integrated into pSpCas9(BB)-2A-GFP (PX458) (Addgene no. 48138) and 2μg of plasmid was electroporated into 10^6^ cells using a Lonza 4D-Nucleofector according to the manufacturer’s protocol for HCT116 cells. GFP-positive cells were sorted 24hrs after electroporation using a Sony SH800 cell sorter with the brightest 5% being pooled for recovery in medium containing 50% FBS for 4 days. Sorted populations were then seeded at low density and individual clones were isolated after 10 days of outgrowth. Individual clones were validated by sequencing.

### Survival Experiments

To generate survival curves for and MEF cell lines, seeding assays were first performed on10^4^ cells allowed to grow for 7 days respectively. Following this, the growth medium was removed and the cells were washed briefly with PBS before the addition of crystal violet stain (0.5% crystal violet in 25% methanol). Cells were stained for 5min, washed with ddH_2_O and dried before scanning. Cell seeding numbers were calculated relative to WT cell density. Survival curves were then generated by seeding 300-400 WT or *Brca1*^+/−^*53bp1*^-/-^ cells or 650-750 *Brca1*^−/−^*53bp1*^-/-^ cells in triplicate for each drug concentration in a 96-well plate. Olaparib was added 24hrs later to the indicated final concentrations.

Survival curves for *BARD1^AID^*^/AID^ and derivative cell lines were generated as previously described^14^. Briefly, 300/1000 (DMSO/IAA condition) *BARD1*^AID/AID^ cells, 300/700 *BARD1*^AID/*AID*^*53BP1*^-/-^ cells or 400/1500 *BARD1*^AID/*AID*^*SHLD3*^-/-^ cells, per well were seeded in the presence of doxycycline (2 μg ml^−1^) in triplicate for each drug concentration in a 96-well plate. After 24hrs, IAA (1 mM) or carrier (DMSO) was added. One hour after IAA addition, olaparib was added to the indicated final concentrations.

For both cell lines; seven days after drug addition, the medium was replaced with phenol red-free DMEM (Thermo Fisher, 21063-029) supplemented with 10% FBS, penicillin– streptomycin, 2mM L-glutamine and 10μg ml^−1^ resazurin (Sigma-Aldrich, R7017). Plates were then returned to the incubator for 2–4hrs or until the growth medium in untreated control wells began to develop a pink colour. Relative fluorescence was measured with a BMG LABTECH CLARIOstar plate reader. The mean of three technical repeats after background subtraction was taken as the value for a biological repeat and three biological repeats were performed for each experiment. All survival curves presented in this Article represent the mean of three biological repeats ± SEM.

### Immunofluorescence

For RAD51 foci quantification in *BARD1^AID^*^/AID^ and derivative cell lines, 2 × 10^5^ cells were passed through a 70-μm mesh cell strainer and seeded on 3 fibronectin-coated glass coverslips (13 mm) in a single well of a 6-well plate in the presence of doxycycline (2μg ml^−1^). After 24hrs, IAA was added to a final concentration of 1 mM. Cells were irradiated (5 Gy) 2hrs after IAA addition and fixed in 2% PFA 2hrs after irradiation. Staining of all fixed cells began with 15 min blocking (3% BSA, 0.1% Triton X-100 in PBS), followed by 1hr incubation with primary antibody in a humidity chamber. For experiments in which cells were treated with EdU, the Click-iT EdU Cell Proliferation Kit, Alexa Fluor 647 (Thermo Fisher, C10340) was used to label EdU-positive cells according to the manufacturer’s protocol between blocking and primary antibody incubation.

The following primary antibodies were used at the indicated concentrations: rabbit anti-RAD51 (1:1,000, 70-001 BioAcademia), mouse anti-γH2AX (1:500, 05-636 Millipore). Following primary, coverslips were washed 3 times with PBS containing 0.1% Triton X-100 before incubation with secondary antibody for 1 h in a humidity chamber. Secondary antibodies used in this study were: goat anti-mouse Alexa Fluor 488 (1:500, A-11001 Invitrogen) and goat anti-rabbit Alexa Fluor 568 (1:500, A-11011 Invitrogen). Coverslips were then washed 3 times with PBS containing 0.1% Triton X-100, once with PBS, and mounted on glass microscope slides using a drop of ProLong Gold antifade reagent with DAPI (Life Technologies, P36935).

Images were acquired on a Leica DMi8 widefield microscope. CellProfiler (Broad Institute) was used for foci quantification. Images were visualized and saved in Fiji and assembled into Fig. in Affinity Designer.

### Cytological analysis

Metaphase spreads were prepared by standard methods. In brief, detached cells were resuspended in KCl 75 mM for 20 min, before being fixed in Carnoy’s fixative. Approximately 20 μl of cell suspension was dropped onto clean slides and left to dry overnight. The cells were then stained with propidium iodide 0.5μg/ml in PBS for 20 min, rinsed and the slides mounted in Vectashield/DAPI (Vector Labs). The slides were analysed blind, and a minimum of 40 metaphases were acquired using an Olympus BX60 microscope for epifluorescence equipped with a Sensys CCD camera (Photometrics). Images were collected using Genus Cytovision software (Leica).

### Statistics

Prism 10 (Graphpad Software) was used for graphing and statistical analysis. Relevant statistical methods for individual experiments are detailed within Fig. legends.

## Extended Data: Titles and legends

**Supplementary Fig. 1: Related to Figure 1.**

**A)** PCR amplicons from genomic DNA obtained by ear biopsies from mice of the indicated genotype. Bands of different size correspond to the *Shld2^tm1a^* allele (274 bp), *Shld2^tm1c^* allele (554 bp), *Shld2^tm1d^* allele (240 bp) and *WT* allele (391 bp).
**B)** PCR amplicons from genomic DNA obtained by ear biopsies from mice of the indicated genotype. Bands of different size correspond to the *Shld3^EM1-B6^* allele (left) or *Shld3^EM2-B6^* allele (right) (∼312 bp) and *WT* allele (459 bp).
**C)** PCR amplicons from genomic DNA obtained by ear biopsies (left), splenic b cells or purified splenic b cells (right) from mice of the indicated genotype. Bands of different size correspond to the *Ctc1^fl^* allele (413 bp), *Ctc1^-/-^* allele (658 bp) and *WT* allele (210 bp).
**D)** Peripheral blood stained as indicated. Reticulocytes (RET), micronucleated reticulocytes (MN-RET), normochromatic erythrocytes (NCE) and micronucleated normochromatic erythrocytes (MN-NCE). Representative data, n ≥ 5 mice of each sex per genotype.
**E)** MN-NCE as a percentage of total NCEs from tail vein blood of mice at 8-12 weeks. n=5-12 mice of each sex per genotype. Significance was determined by unpaired two tailed *t*-test, Mean ± SEM.
**F)** Flow cytometric sub-classification of nature splenic b cell fractions. n = 4-8 mice per genotype. Significance was determined by unpaired two tailed *t*-test, Mean±SEM.
**G)** Absolute numbers of B220^+^ B cells in the bone marrow (one femur and one tibia) and spleen and live lymphocytes in the thymus. n = 10-18 mice per genotype. Significance was determined by unpaired two tailed *t*-test, Mean ± SEM.
**H)** Absolute numbers of B cell precursors in the bone marrow (one femur and one tibia). n = 15-19 mice per genotype. Significance was determined by unpaired two tailed *t*-test, Mean±SEM.
**I)** Flow cytometric sub-classification of mature splenic b cell fractions. n = 6-17 mice per genotype. Significance was determined by unpaired two tailed *t*-test, Mean ± SEM.
**J)** Absolute numbers of thymic T-cells, n = 9-17 mice per genotype.
**K)** Absolute numbers of T cell precursors in double negative (CD4-CD8-) in the thymus, n = 9-17 mice per genotype.
**L)** Flow cytometry analysis of B cell development in the bone marrow of *Mb1^+/Cre^*, *Shld2^-/- or^ Ctc1^fl/fl^ Mb1^+/Cre^ mice*; gating on B220^+^CD43^+^ (left, Hardy fractions A, B and C) and on B220^+^CD43^-^ (right, Hardy fractions D, E and F). Representative data; *n* = 3 experiments.
**M)** Flow cytometry analysis of mature b cell populations in the spleen of *Mb1^+/Cre^*, *Shld2^-/- or^ Ctc1^fl/fl^ Mb1^+/Cre^ mice*; gating on B220^+^CD19^+^. Representative data; *n* = 3 experiments.
**N)** Flow cytometry analysis of B cell development in the bone marrow of WT, *53bp1*^-/-^, *Shld2/3^-/-^ mice*; gating on B220^+^CD43^+^ (top, Hardy fractions A, B and C) and on B220^+^CD43^-^ (bottom, Hardy fractions D, E and F). Representative data; *n* > 8 experiments.
**O)** Flow cytometry analysis of mature b cell populations in the spleen of WT, *53bp1*^-/-^, *Shld2/3^-/-^ mice*; gating on B220^+^CD19^+^. Representative data; *n* > 8 experiments.
**P)** Flow cytometry analysis of T cell development in the thymus of WT, *53bp1*^-/-^, *Shld2/3^-/-^ mice*; gating on live cells (top) and on CD8^-^ CD4^-^ (bottom, double negative populations). Representative data; *n* >3 experiments.
**Q)** CTV-labelled purified B cells were stimulated as indicated and stained for surface IgG/IgE on day 4. Representative of *n* > 6 experiments.
**R)** Splenic B cells cultured with the indicated stimuli (96 h) and stained for surface IgG1, IgE, IgG2b or IgG3. *n* = 11-14 mice per genotype. CSR 100%, mean immunoglobulin isotype switch frequency of 2 control animals in each experiment. Significance was determined by two-way ANOVA with Tukey’s correction. Mean ± SEM.

**Supplementary Fig. 2: Related to Figure 2.**

**A)** IgM-to-IgA CSR frequencies in *Shld3* knockout CH12-F3 cell lines, *n* = 3 independent experiments. Mean ± SEM. Representative flow cytometry plots for IgM-to-IgA CSR. Representative data, *n* > 3 independent experiments.
**B)** IgM-to-IgA CSR frequencies in *Shld1* knockout CH12-F3 cell lines, *n* = 3 independent experiments. Mean ± SEM. Representative flow cytometry plots for IgM-to-IgA CSR. Representative data, *n* > 3 independent experiments.
**C)** B cell *Shld1* interactome as defined by LC–MS/MS and LFQ. Scatter plot depicts log2 fold-enrichment of indicated TwinStep–immunocomplexes across 2 independent experiments.
**D)** B cell *Shld1* IR-dependent interactome as defined by LC–MS/MS and LFQ. Scatter plot depicts log2 fold-enrichment of indicated TwinStep–immunocomplexes across 2 independent experiments.
**E)** Western blot analysis of CH12-F3 cell extracts from pulldown experiments. Cells were pre-treated with ATM, ATR inhibitor or both for 1hr prior to irradiation. Representative of n = 2 independent experiments.
**F)** Western blot analysis of CH12-F3 cell extracts from pulldown experiments. Representative of n = 2 independent experiments.
**G)** Splenic B cells cultured with the indicated stimuli (96 h) and stained for surface IgG1, IgE, IgG2b or IgG3. *n* = 3-5 mice per genotype. CSR 100%, mean immunoglobulin isotype switch frequency of 2 control animals in each experiment. Mean ± SEM.
**H)** CTV-labelled purified B cells were stimulated as indicated and stained for surface IgG on day 4. Representative of *n* = 2 experiments.

**Supplementary Fig. 3: Related to Figure 3.**

**A)** PCR amplicons from genomic DNA obtained by ear biopsies (left), splenic b cells or purified splenic b cells (right) from mice of the indicated genotype. Bands of different size correspond to the *Rev3l^fl^* allele (458 bp), *Rev3l^-/-^* allele (300 bp) and *WT* allele (384 bp).
**B)** CTV-labelled purified B cells were stimulated as indicated and stained for surface IgG/IgE on day 4. Representative of *n* > 5 experiments.

**Supplementary Fig. 4: Related to Figure 4.**

**A)** Flow cytometric sub-classification of nature splenic b cell fractions. n = 4-6 mice per genotype. Significance was determined by unpaired two tailed *t*-test, Mean±SEM.
**B)** Flow cytometry analysis of B cell development in the bone marrow of *Mb1^+/Cre^*, *Rev7^f/lf^ Mb1^+/Cre or^ Rev3l^fl/fl^ Mb1^+/Cre^ mice*; gating on B220^+^CD43^+^ (top, Hardy fractions A, B and C) and on B220^+^CD43^-^ (bottom, Hardy fractions D, E and F). Representative data; *n* = 3 experiments.
**C)** Flow cytometry analysis of mature b cell populations in the spleen of *Mb1^+/Cre^*, *Rev7^f/lf^ Mb1^+/Cre or^ Rev3l^fl/fl^ Mb1^+/Cre^ mice*; gating on B220^+^CD19^+^. Representative data; *n* = 3 experiments.
**D)** PCR amplicons from genomic DNA obtained by ear biopsies (left), splenic b cells or purified splenic b cells (right) from mice of the indicated genotype. Bands of different size correspond to the *p53^fl^* allele (353 bp), *p53^-/-^* allele (450 bp) and *WT* allele (200 bp).
**E)** Flow cytometric sub-classification of nature splenic b cell fractions. n =6-12 mice per genotype. Significance was determined by unpaired two tailed *t*-test, Mean±SEM.
**F)** Flow cytometry analysis of B cell development in the bone marrow of *Rev^+/+^ p53^fl/fl^ Mb1^+/Cre or^ Rev3l^fl/fl^ p53^fl/fl^ Mb1^+/Cre^ mice*; gating on B220^+^CD43^+^ (top, Hardy fractions A, B and C) and on B220^+^CD43^-^ (bottom, Hardy fractions D, E and F). Representative data; *n* = 3 experiments.
**G)** Flow cytometry analysis of mature b cell populations in the spleen of *Rev^+/+^ p53^fl/fl^ Mb1^+/Cre or^ Rev3l^fl/fl^ p53^fl/fl^ Mb1^+/Cre^ mice*; gating on B220^+^CD19^+^. Representative data; *n* = 3 experiments.
**H)** Flow cytometry analysis of B cell development in the bone marrow of MD4^+/Tg^, Rev3^fl/fl^ Mb1^+/Cre^ MD4^+/Tg^, 53BP1^-/-^ MD4^+/TG^ mice; gating on B220^+^. Representative data; *n* = 3 experiments.

**Supplementary Fig. 5: Related to Figure 5.**

**A)** Survival of the indicated BARD^AID/AID^ cell lines grown for 7 days in the presence of doxycycline (2 μg ml^-1^) and the indicated doses of olaparib, n=3 biological experiments, Mean±SEM.
**B)** Quantification of between 40 and 65 metaphases analysed per genotype/experiment. Significance was determined by two-sided Kruskal–Wallis *H* test with Dunn’s correction for multiple comparisons. ****P<0.0001, ***P<0.0002, **P<0.002, *P<0.05. All comparisons not indicated in are not significant.

