## Supplementary figures and images for "Shieldin and CST co-orchestrate DNA polymerase-dependent tailed-end joining reactions independently of 53BP1-governed repair pathway choice"

### Extended Data Figure 1

#### Extended Data Fig. 1:

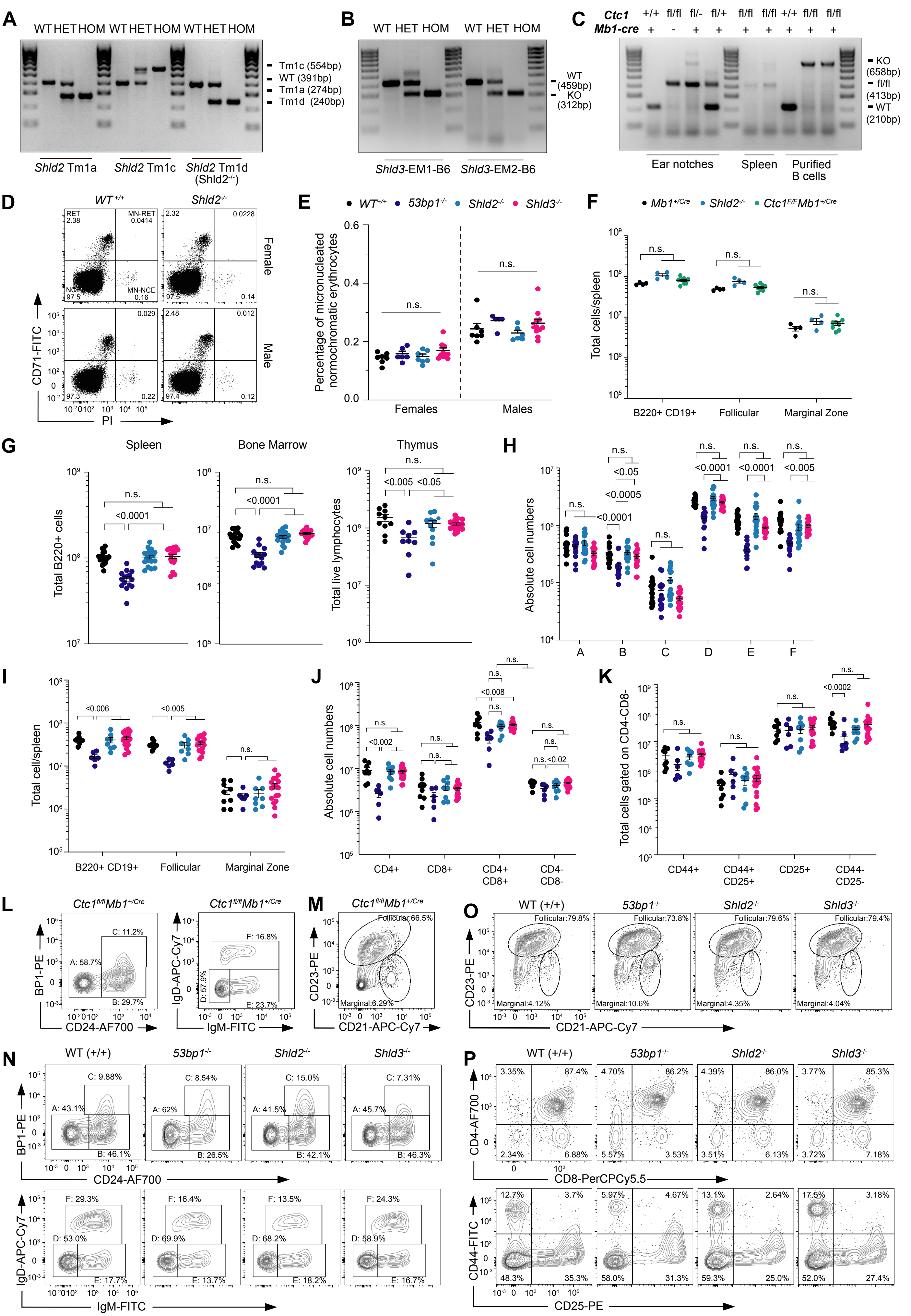

##### Extended Data Fig. 1 Continued:

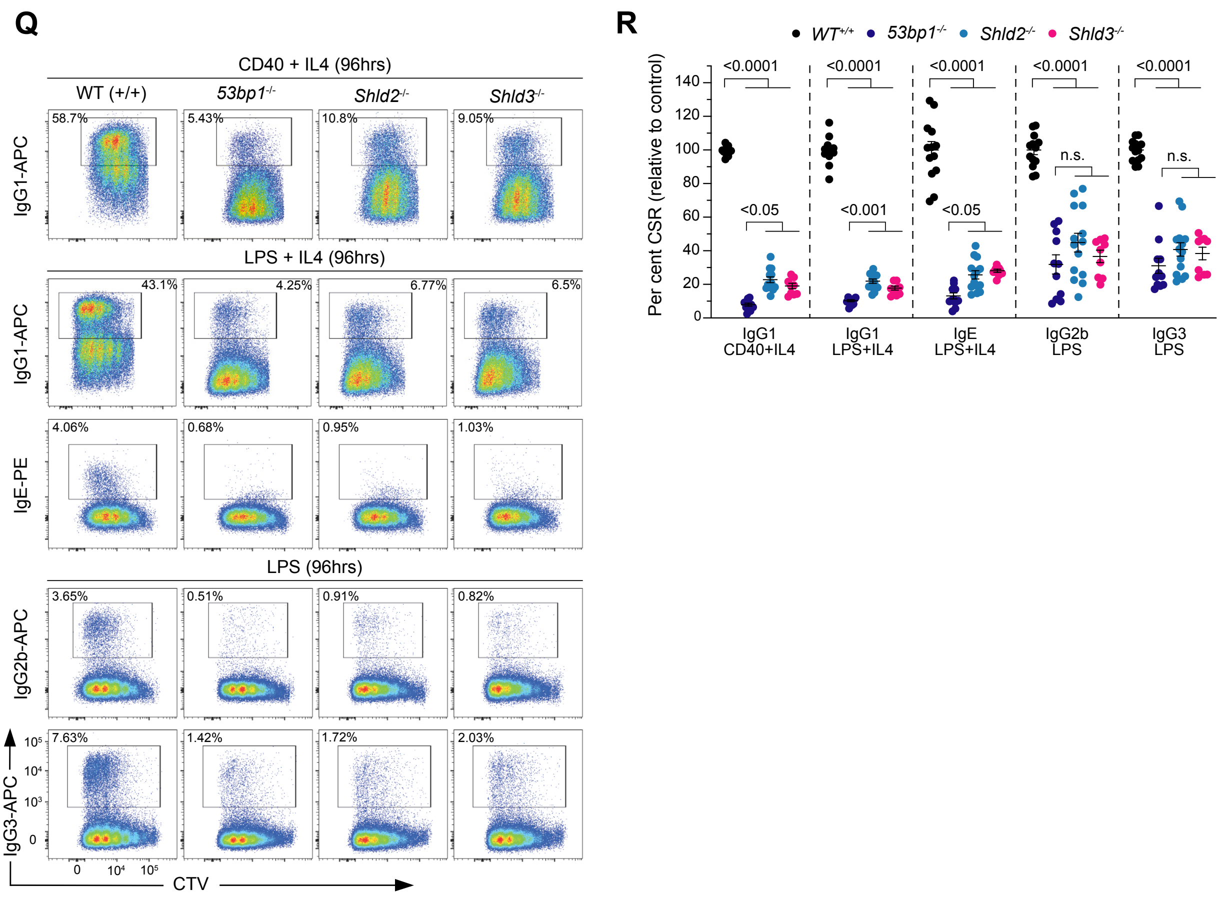

### Extended Data Figure 2

##### Extended Data Fig. 2:

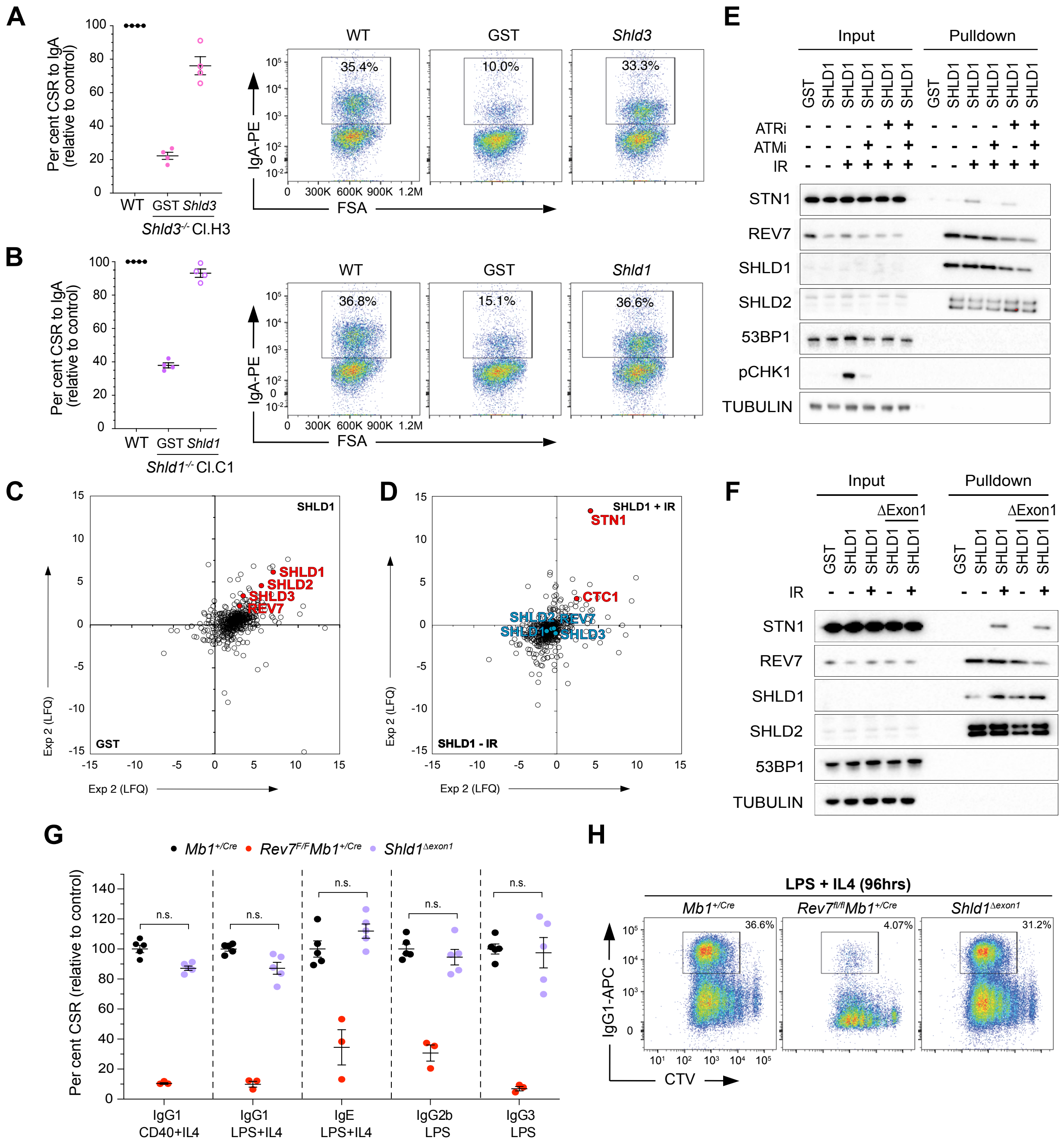

### Extended Data Figure 3

#### Extended Data Fig. 3:

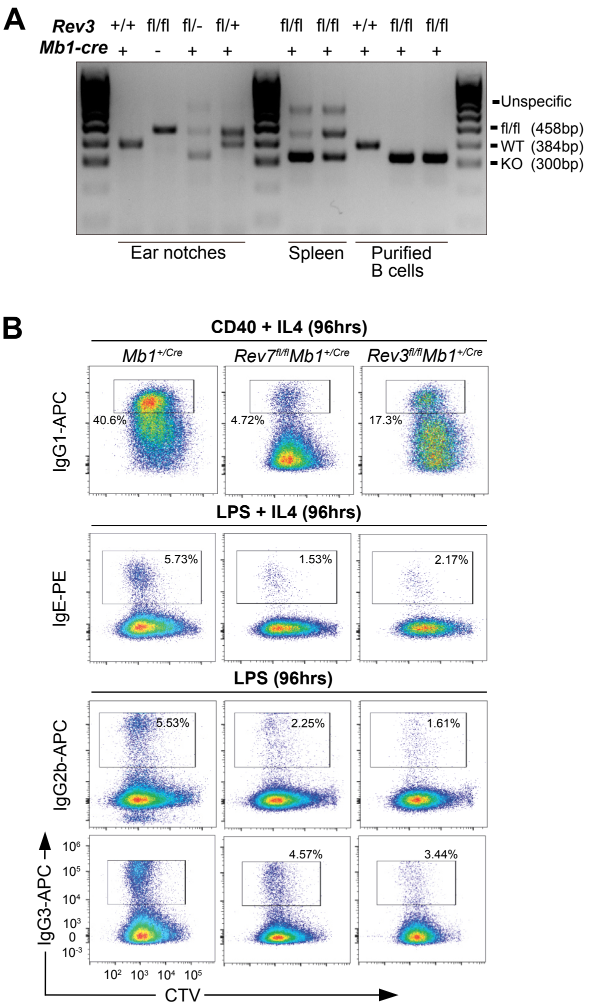

### Extended Data Figure 4

##### Extended Data Fig. 4:

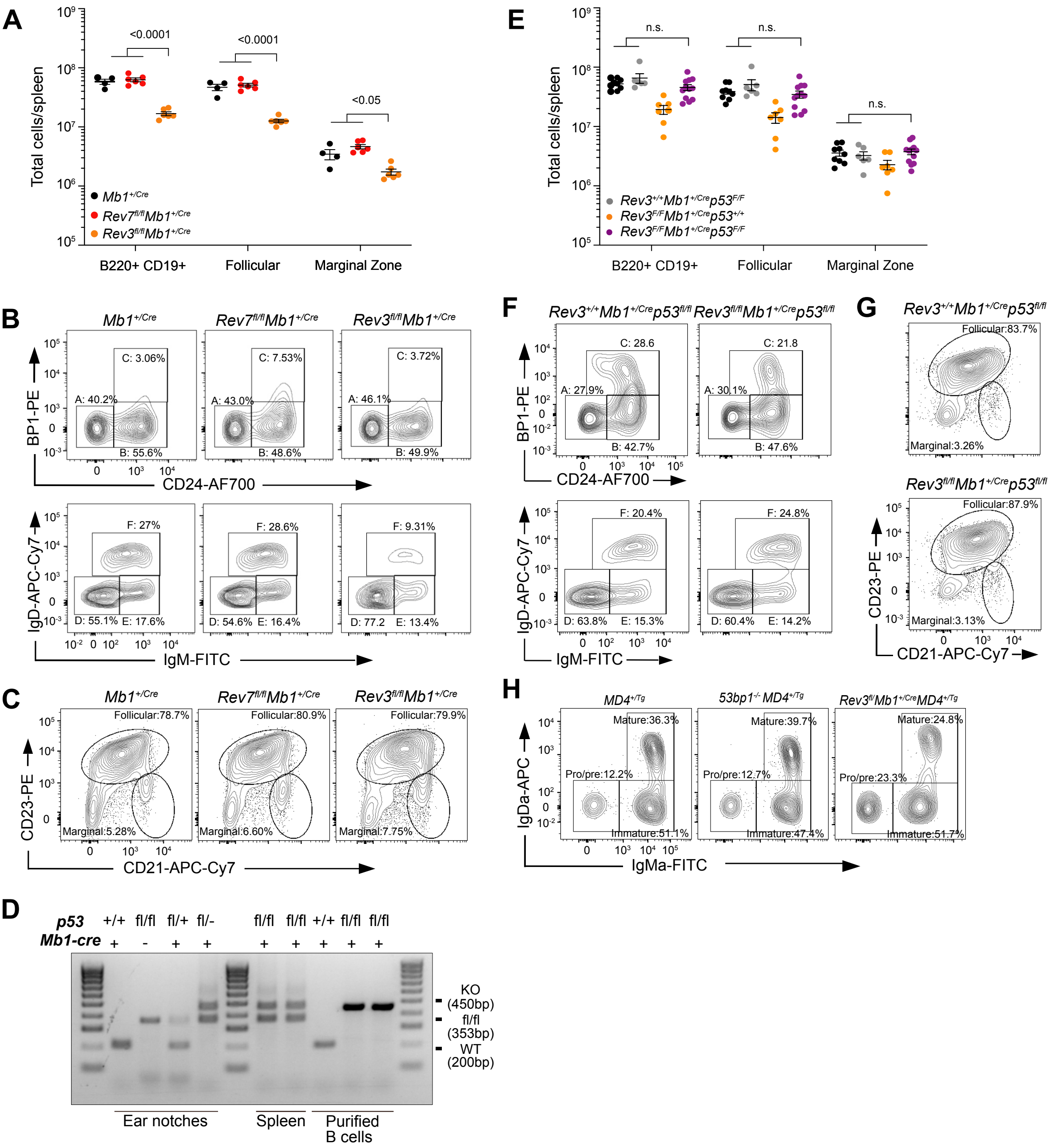

### Extended Data Figure 5

**Extended Data Fig. 5:**

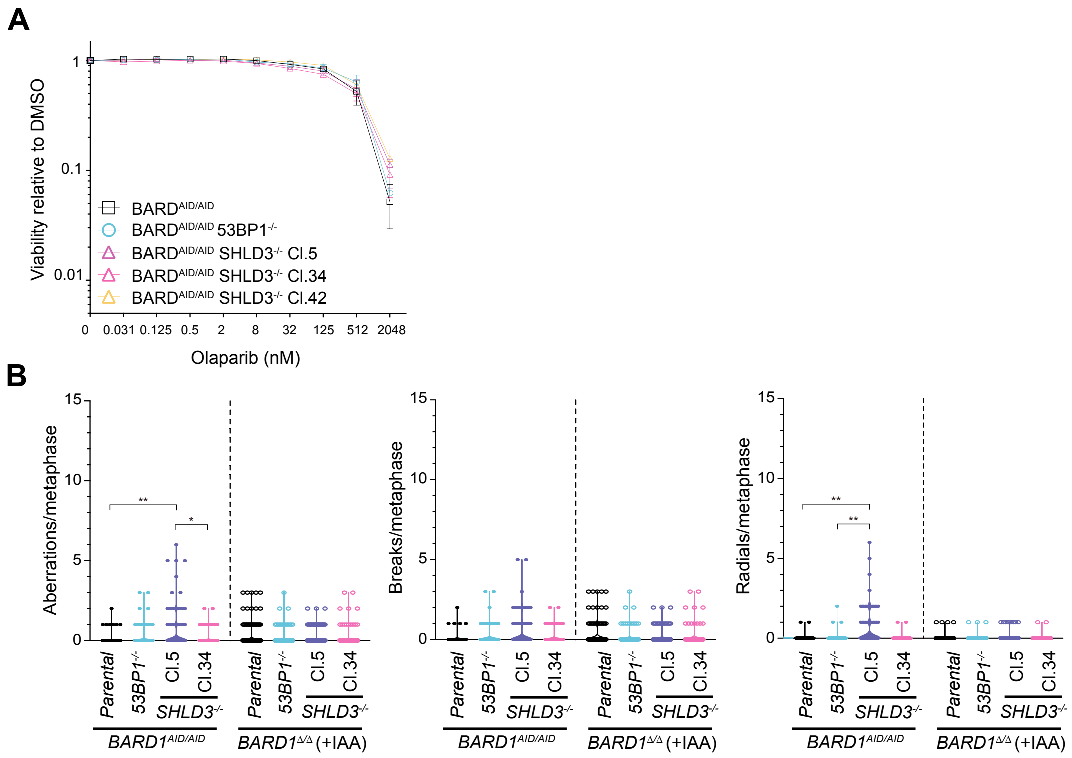
